# Context-Dependent Modification of PFKFB3 in Hematopoietic Stem Cells Promotes Anaerobic Glycolysis and Ensures Stress Hematopoiesis

**DOI:** 10.1101/2023.03.16.532898

**Authors:** Shintaro Watanuki, Hiroshi Kobayashi, Yuki Sugiura, Masamichi Yamamoto, Daiki Karigane, Kohei Shiroshita, Yuriko Sorimachi, Shinya Fujita, Takayuki Morikawa, Shuhei Koide, Motohiko Oshima, Akira Nishiyama, Koichi Murakami, Miho Haraguchi, Shinpei Tamaki, Takehiro Yamamoto, Tomohiro Yabushita, Yosuke Tanaka, Go Nagamatsu, Hiroaki Honda, Shinichiro Okamoto, Nobuhito Goda, Tomohiko Tamura, Ayako Nakamura-Ishizu, Makoto Suematsu, Atsushi Iwama, Toshio Suda, Keiyo Takubo

**Author notes:** These authors contributed equally (Shintaro Watanuki, Hiroshi Kobayashi). Corresponding Authors: Keiyo Takubo, M.D., Ph.D. and Hiroshi Kobayashi, M.D., Ph.D. Department of Stem Cell Biology, Research Institute National Center for Global Health and Medicine 1-21-1 Toyama, Shinjuku-ku, Tokyo 162-8655, Japan / (KT) | (HK), Yuki Sugiura, Ph.D. Center for Cancer Immunotherapy and Immunobiology Kyoto University Graduate School of Medicine Yoshida-Konoe, Sakyo-ku, Kyoto 606-8501, Japan. Scientific category: Hematopoiesis and Stem Cells.

## Abstract

Metabolic pathways are plastic and rapidly change in response to stress or perturbation. Current metabolic profiling techniques require lysis of many cells, complicating the tracking of metabolic changes over time after stress in rare cells such as hematopoietic stem cells (HSCs). Here, we aimed to identify the key metabolic enzymes that define differences in glycolytic metabolism between steady-state and stress conditions in HSCs and elucidate their regulatory mechanisms. Through quantitative ^13^C metabolic flux analysis of glucose metabolism using high-sensitivity glucose tracing and mathematical modeling, we found that HSCs activate the glycolytic rate-limiting enzyme phosphofructokinase (PFK) during proliferation and oxidative phosphorylation (OXPHOS) inhibition. Real-time measurement of adenosine triphosphate (ATP) levels in single HSCs demonstrated that proliferative stress or OXPHOS inhibition led to accelerated glycolysis via increased activity of PFKFB3, the enzyme regulating an allosteric PFK activator, within seconds to meet ATP requirements. Furthermore, varying stresses differentially activated PFKFB3 via PRMT1-dependent methylation during proliferative stress and via AMPK-dependent phosphorylation during OXPHOS inhibition. Overexpression of *Pfkfb3* induced HSC proliferation and promoted differentiated cell production, whereas inhibition or loss of *Pfkfb3* suppressed them. This study reveals the flexible and multilayered regulation of HSC glycolytic metabolism to sustain hematopoiesis under stress and provides techniques to better understand the physiological metabolism of rare hematopoietic cells.

**Key Points:**

- Combined isotope tracing, mathematical modeling, and single cell ATP analysis enable high-resolution evaluation of blood cell metabolism.
- Under stress, HSCs quickly accelerate glycolysis to meet ATP demands and maintain hematopoiesis via context-dependent PFKFB3 activation.

## Introduction

Activities governing nutrient requirements and metabolic pathways in individual cells maintain tissue homeostasis and respond to stress through metabolite production. Adenosine triphosphate (ATP), produced via cytosolic glycolysis and mitochondrial oxidative phosphorylation (OXPHOS), is the universal energy currency of all organisms; it regulates all anabolic or catabolic cellular activities^1–3^. Precise control of intracellular ATP concentrations is critical, as ATP is the rate determiner of many ATP-dependent biochemical reactions^4–9^.

Hematopoietic stem cells (HSCs) are tissue stem cells at the apex of the hematopoietic hierarchy; their function is maintained throughout life by a rigorous metabolic program and a complex interplay of gene expression, epigenetic regulation, intracellular signaling, chromatin remodeling, autophagy, and environmental factors^10–14^. Conventional analyses of the metabolic programs of hematopoietic stem and progenitor cells (HSPCs) have revealed diverse differentiation potentials and cell-cycling statuses and coordinated activities that maintain hematopoiesis^15–26^. Among the HSPC fractions, HSCs possess unique cell cycle quiescence, high self-renewal and differentiation capacity in response to stimuli, and resistance to cellular stress, including reactive oxygen species and physiological aging^10,27–30^. These properties are regulated by a balance between glycolysis and mitochondrial OXPHOS, requiring biosynthesis of ATP and various macromolecules that confer resilience to stress^31^. Among the known regulators of ATP-producing pathways, glycolytic enzymes maintain HSCs and hematopoietic progenitor cells (HPCs) by regulating cellular survival and cell cycle quiescence^32–34^. Loss of mitochondrial genes in HPSCs also induces HSC differentiation defects^35–37^. Moreover, disrupting the mitochondrial complex III subunit depletes both differentiated hematopoietic cells and quiescent HSCs^21^. Although glycolysis and the tricarboxylic acid (TCA) cycle are metabolically linked, pyruvate dehydrogenase kinase activity, which can uncouple these pathways, is required to maintain HSC function^32,38^.

During HSC division, cell metabolism is reprogrammed to activate fatty acid β-oxidation (FAO) and purine metabolism^39–41^. Furthermore, Liang et al. reported that activated HSCs mainly rely on glycolysis as their energy source^42^. However, the mechanisms by which each ATP-producing pathway and their connections are differentially regulated between HSCs and differentiated cells at steady state, during cell cycling, or during stress remain unknown. Recently, it has been shown that deeply quiescent HSCs do not activate cell cycle under stress^43–45^. Therefore, it remains unclear whether metabolic changes such as the individual ATP-producing pathways and their interconnections occur uniformly in all HSCs, including these deeply quiescent HSCs. Furthermore, the underlying hub metabolic enzyme responsible for changes in the metabolic system of HSCs under stress has not been identified. HSCs are essential for cell therapy, including HSC transplantation, and in order to comprehensively elucidate the metabolic systems that have attracted attention as their regulatory mechanisms, recent studies have included metabolomic analyses using rare cell types such as HSCs^23,46–49^, as well as isotope tracer analyses of undifferentiated hematopoietic cells purified after *in vivo* administration of isotopic glucose^50^. Although these approaches are useful for obtaining comprehensive information on intracellular metabolites, they are not suited to track real-time changes in cellular metabolism at high resolution. Therefore, new approaches are necessary to analyze metabolites quantitatively and continuously without disturbing the physiological states of single cells while integrating the recently reported metabolome analysis techniques. In this study, we aimed to identify the key metabolic enzymes that define differences in glycolytic metabolism between steady-state and stress conditions in HSCs and elucidate their regulatory mechanisms using a quantitative and mathematical approach. Our findings provide a platform for quantitative metabolic analysis of rare cells such as HSCs, characterize the overall metabolic reprogramming of HSCs during stress loading, and highlight the key enzyme involved in this process.

### Data sharing

All relevant data are available from the corresponding author upon reasonable request.

## Results

### HSC cell cycling increases anaerobic glycolytic flux

To determine how cell cycle progression alters HSC metabolism *in vivo*, we intraperitoneally and intravenously treated mice with 5-fluorouracil (5-FU) to induce HSC cell cycling (Supplemental Figure 1A). For analysis after 5-FU administration, the Lineage (Lin)^-^ Sca-1^+^ c-Kit^+^ (LSK) gate was expanded to include HSCs with decreased c-Kit expression levels early after 5-FU treatment, for example high Sca-1-expressing cells and c-Kit-high to -dim Lin^-^ cells, based on the previous report^51^ ^41^(Supplemental Figure 1B). This expanded LSK gate was consistent with the patterns of c-Kit expression observed in endothelial protein C receptor (EPCR)^+^ Lin^-^ CD150^+^ CD48^-^ cells (Supplemental Figure 1C) with high stem cell activity after 5-FU administration^41^. We observed a transient decrease in the number of quiescent HSCs (Ki67^-^) and an increase in the number of cell-cycling HSCs (Ki67^+^) on day 6 after 5-FU treatment (Supplemental Figure 1D). Along with the loss of cell quiescence, ATP concentration in HSCs decreased transiently on day 6 (Supplemental Figure 1E). Because the route of administration of 5-FU (intraperitoneal or intravenous) made no difference in the Ki67 positivity rate of HSCs (Supplemental Figure 1F), we administered 5-FU intraperitoneally for remaining experiments. Two methods were used to test whether cell cycle progression of HSCs after 5-FU treatment depends on the expression of EPCR. First, phosphorylation of Rb (pRb), a marker of cell cycle progression^52^, was analyzed in HSCs after 5-FU treatment. Analysis of EPCR^+^ and EPCR^-^ HSCs showed increased pRb in HSCs from 5-FU-treated mice in both fractions compared to HSCs from phosphate-buffered saline (PBS)-treated mice, regardless of EPCR expression (Supplemental Figure 1G-H). Second, we used a G_0_ marker mouse line^53^. These mice expressed a fusion protein of the p27 inactivation mutant p27K^-^ and the fluorescent protein mVenus (G_0_ marker), allowing prospective identification of G_0_ cells. We tested whether the expression of G_0_ marker in HSCs was altered after 5-FU administration to the G_0_ marker mice (Supplemental Figure 1I) and found that 5-FU treatment reduced the frequency of G_0_ marker-positive HSCs, regardless of the EPCR expression (Supplemental Figure 1J-K). This was not observed in the PBS group. These results indicated that 5-FU administration induced cell cycle progression of entire HSCs in mice.

HSC cell cycling is preceded by the activation of intracellular ATP-related pathways that metabolize extracellular nutrients, including glucose^39,40^, which are utilized in both ATP-producing and -consuming pathways, determining cellular ATP levels. Therefore, we examined the metabolic flux of glucose by performing *in vitro* IC-MS tracer analysis with uniformly carbon-labeled (U-^13^C_6_) glucose to determine the pathways driving changes in ATP in 5-FU-treated HSCs (Figure 1A; Table S2). To avoid metabolite changes, samples were continuously chilled on ice during cell preparation, and the process from euthanasia to cell preparation was performed in the shortest possible time (see “**Preparation and storage of *in vitro* U-^13^C_6_-glucose tracer samples**” section under "**Methods**" for more information). We found that changes in metabolite levels before and after sorting were present but limited (Supplemental Figure 2A). This result is consistent with the finding that the cell purification process does not significantly affect metabolite levels when sufficient care is taken in cell preparation^50^. In 5-FU-treated HSCs, the levels of glycolytic metabolites derived from U-^13^C_6_-glucose were double those observed in PBS-treated HSCs (Figure 1B-C; Supplemental Figure 2B). The total levels of TCA cycle intermediates derived from U-^13^C_6_-glucose were similar between PBS- and 5-FU-treated cells (Figure 1D; Supplemental Figure 2B). Levels of U-^13^C_6_-glucose-derived intermediates involved in the pentose phosphate pathway (PPP) and nucleic acid synthesis (NAS) were two-fold higher in 5-FU-treated than in PBS-treated HSCs, whereas no significant differences in the levels of metabolites were observed between both groups (Figure 1E-F; Supplemental Figure 2B). Notably, the labeling rate of metabolites during the first half of glycolysis was almost 100% in both groups, allowing us to easily track the labeled metabolites (Supplemental Figure 2C-E).This was thought to be due to the rapid replacement of unlabeled metabolites with labeled metabolites during exposure to U-^13^C_6_-glucose because of the generally rapid glycolytic reaction. Conversely, the labeling rate of TCA cycle intermediates was consistently lower than that of glycolysis and PPP (Supplemental Figure 2D), suggesting that PBS- and 5-FU-treated HSCs prefer anaerobic glycolysis over aerobic glycolysis. To directly compare the metabolic systems of PBS- or 5-FU-treated HSCs, we conducted a Mito stress test using a Seahorse flux analyzer. Compared to PBS-treated HSCs, 5-FU-treated HSCs exhibited a higher extracellular acidification rate (ECAR), while their oxygen consumption rate (OCR) remained equal to that of PBS-treated HSCs (Figure 1G-H; Supplemental Figure 3A-B). After oligomycin treatment, PBS- and 5-FU-treated HSCs showed an increase in ECAR, suggesting a flexible activation of glycolysis upon OXPHOS inhibition (Figure 1G; Supplemental Figure 3A). Meanwhile, a decrease in OCR was more clearly observed in the 5-FU-treated HSCs (Figure 1H; Supplemental Figure 3B). Next, we evaluated whether glucose uptake in HSCs after 5-FU administration was differentially affected by the expression of EPCR. The fluorescent analog of glucose, 2-(N-(7-nitrobenz-2-oxa-1,3-diazol-4-yl)amino)-2-deoxyglucose (2-NBDG), was administered intravenously to mice^50^ and its uptake in EPCR^+^ and EPCR^-^ HSCs was assayed (Figure 1I). Regardless of the EPCR expression, the 2-NBDG uptake was greater in HSCs treated with 5-FU than in those treated with PBS (Figure 1J-L). Increased 2-NBDG uptake in 5-FU-treated HSCs was also observed in an *in vitro* 2-NBDG assay (Supplemental Figure 1L). Notably, even in the PBS-treated group, HSCs with high NBDG uptake were more proliferative than those with low NBDG uptake, similar to the state of HSCs after 5-FU administration (Supplemental Figure 1M). After 5-FU administration, there was an overall shift of the population from the G_0_ to G_1_ phase and a correlation between NBDG uptake and cell cycle progression was also observed (Supplemental Figure 1M). In both PBS- and 5-FU-treated groups, the marked variation in glucose utilization depending on the cell cycle suggests a direct link between HSC proliferation and increased glycolytic activity. Furthermore, compared to HSCs cultured under the quiescence-maintaining conditions of HSC achieved by hypoxia, abundant fatty acids, and low cytokines as we previously reported ^54^, HSCs cultured under cytokine-rich proliferative conditions were more resistant to the inhibition of OXPHOS by oligomycin (Supplementary Fig. 1N; Table S1). Overall, the results showed that 5-FU-treated HSCs exhibited activated glycolytic flux, increasing the turnover of ATP. Moreover, glycolytic flux into mitochondria was equally unchanged in PBS- and 5-FU-treated-HSCs, supporting that 5-FU activated anaerobic glycolysis in HSCs.

**Figure 1.**
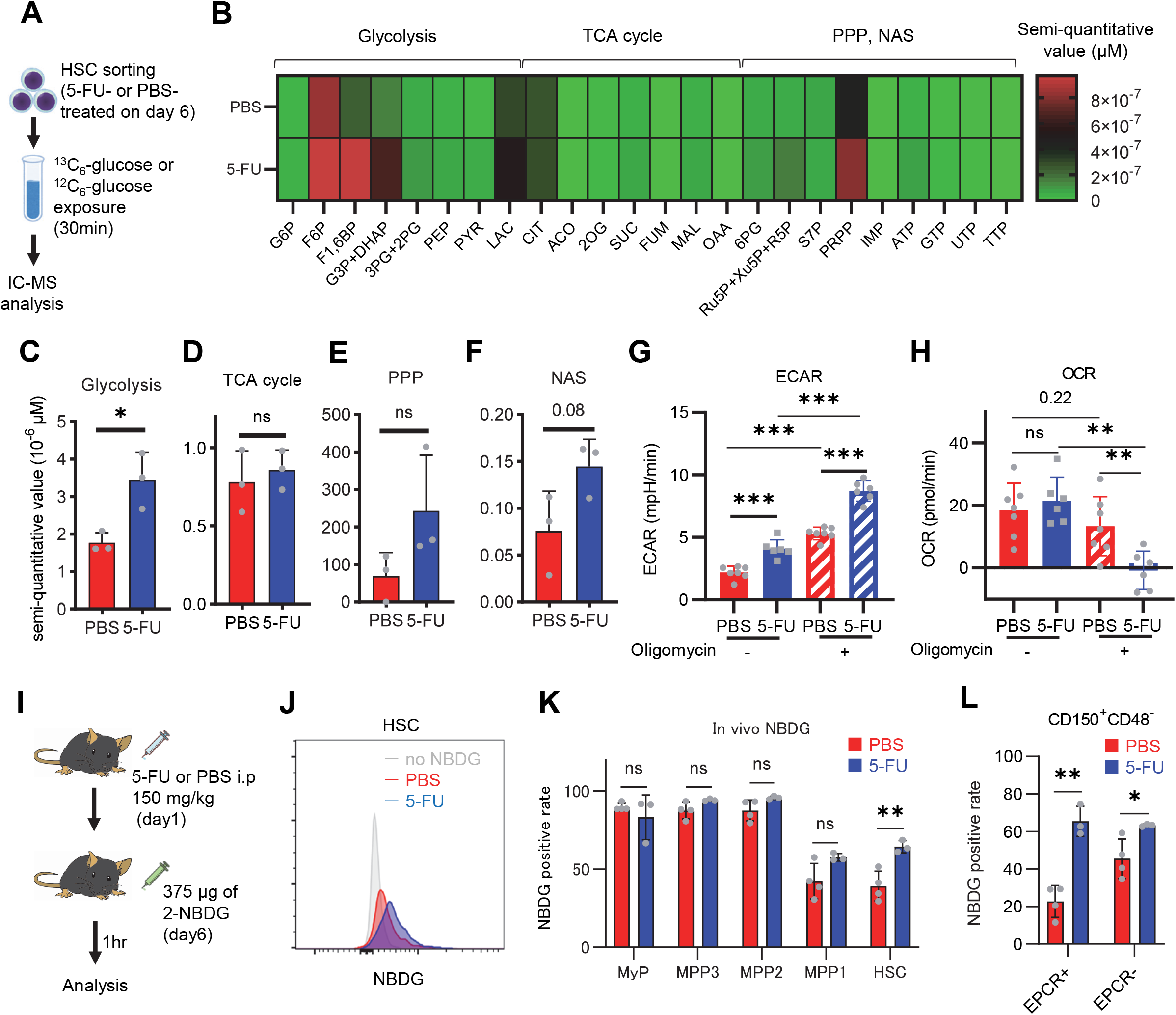
HSC cell cycling increases overall glycolytic flux, but not flux into mitochondria. **(A)** Experimental design used for glucose isotope tracer analysis in HSCs from 5-FU- or PBS-treated mice. **(B)** Heat map of metabolite levels in HSCs derived from mice treated with PBS or 5-FU. **(C-F)** The semi-quantitative value (10^-6^ µM) of U-^13^C_6_-glucose-derived metabolites in glycolysis (C), the first round of TCA cycle (D), the PPP, and nucleotide synthesis (F) in HSCs from 5-FU- or PBS-treated mice (PBS group = 1.0); In (B)-(F), biological replicates from the PBS and 5-FU groups, obtained on three separate days, were pooled, analyzed by IC-MS, quantified based on calibration curve data for each metabolite (see “**Ion chromatography mass spectrometry (IC-MS) analysis**” section in “**Methods**” for details). (**G–H**) A Mito Stress test with the Seahorse flux analyzer on HSCs derived from mice treated with PBS or 5-FU; ECAR (G) and OCR (H) before and after oligomycin treatment. (Data were obtained from n = 7 technical replicates for PBS-treated HSCs and n = 6 for 5-FU-treated HSCs.) **(I)** Experimental schema of *in vivo* 2-NBDG analysis. **(J)** Representative histograms of 2-NBDG analysis (gray: no 2-NBDG, red: PBS group, blue: 5-FU group). **(K)** 2-NBDG positivity in each fraction; data represent four pooled biological replicates for the PBS group and three for the 5-FU group; MyP: myeloid progenitor. **(L)** EPCR expression and 2-NBDG positivity within HSC fractions. Data were extracted from each individual in (K). Data are presented as mean ± SD. * p ≤ 0.05, ** p ≤ 0.01, *** p ≤ 0.001 as determined by Student’s *t*-test (C–F, G–H when comparing the PBS and 5-FU groups, and K–L) or paired-samples t-test (G– H when comparing the conditions before and after exposure to oligomycin within the PBS/5-FU group). Abbreviations: G6P, glucose-6-phosphate; F6P, fructose-6-phosphate; F1,6BP, fructose-1,6-bisphosphate; G3P, glycerol-3-phosphate; DHAP, dihydroxyacetone phosphate; 3PG, 3-phosphoglycerate; 2PG, 2-phosphoglycerate; PEP, phosphoenolpyruvate; PYR, pyruvate; LAC, lactate; Ac-CoA; acetyl-CoA; CIT, citrate; ACO, cis-aconitic acid, isocitrate; 2OG, 2-oxoglutarate; SUC, succinate; FUM, fumarate; MAL, malate; OAA, oxaloacetate; 6PG, glucose-6-phosphate; Ru5P, ribulose-5-phosphate; Xu5P, xylulose-5-phosphate; R5P, ribose-5-phosphate; S7P, sedoheptulose-7-phosphate; E4P, erythrose-4-phosphate; PRPP, phosphoribosyl pyrophosphate; IMP, inosine monophosphate; ATP, adenosine triphosphate; GTP, guanine triphosphate; UMP, uridine monophosphate; UTP, uridine triphosphate; TTP, thymidine triphosphate. See also Fig. S1-3.

### OXPHOS-inhibited HSCs exhibit compensatory glycolytic flux

Previous studies using mouse models of mitochondrial disease or defects in genes involved in electron transport chain and OXPHOS suggest that mitochondrial energy production is essential for maintaining HSC function^21,35–37^, as is the glycolytic system. However, there have been no quantitative reports on how OXPHOS-inhibited HSCs can adapt their metabolism. To understand HSC metabolism under OXPHOS inhibition, we performed *in vitro* U-^13^C_6_-glucose tracer analysis of oligomycin-treated HSCs (Figure 2A; Table S3). Similar to 5-FU-treated HSCs (Figure 1), oligomycin-treated HSCs exhibited glycolytic system activation (Figure 2B-C; Supplemental Figure 2B). Metabolite flux to the TCA cycle and PPP was unchanged, but flux to the NAS was significantly reduced in oligomycin-treated HSCs compared to that in steady-state HSCs (Figure 2D-F; Supplemental Figure 2B). The results suggested that OXPHOS-inhibited HSCs activated compensatory glycolytic flux and suppressed NAS flux. As with 5-FU-treated HSCs, analysis of oligomycin-treated HSCs also showed almost 100% labeling of metabolites in the first half of glycolysis (Supplemental Figure 2F-H), allowing us to easily track the labeled metabolites. To further validate the compensatory glycolytic activation of HSCs under OXPHOS inhibition, a Mito Stress test was performed on HSCs and other differentiated myeloid progenitors (MyPs, Lin^-^Sca-1^-^c-Kit^+^ (LKS^-^) cells). The results showed that ECAR were elevated in HSCs after oligomycin treatment compared to before oligomycin treatment (Figure 2G; Supplemental Figure 3C). No increase in ECAR was observed in MyPs (Figure 2G; Supplemental Figure 3C), supporting that inhibition of OXPHOS activated anaerobic glycolysis specifically in HSCs. Meanwhile, in HSCs, the decrease in OCR after oligomycin administration was less evident compared to MyPs (Figure 2H; Supplemental Figure 3D). In MyPs, both ECAR and OCR were downregulated (Figure 2G-H; Supplemental Figure 3C-D).

**Figure 2.**
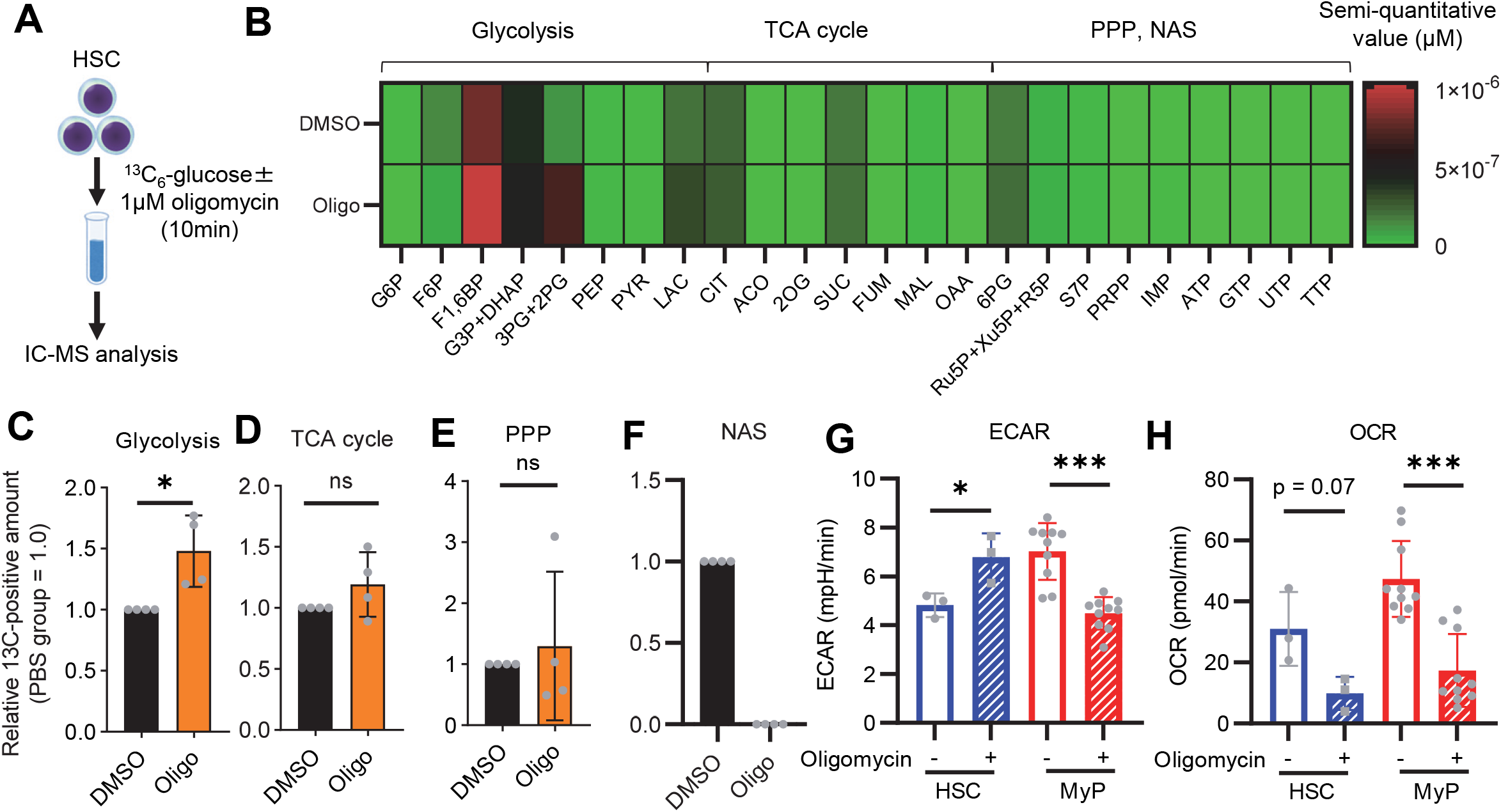
OXPHOS inhibition activates compensatory glycolysis in HSCs. **(A)** Experimental design used for glucose isotope tracer analysis in HSCs treated with the OXPHOS inhibitor oligomycin. **(B)** Heat map of metabolite levels detected by *in vitro* tracer analysis of U-^13^C_6_-glucose in HSCs treated with DMSO or oligomycin (Oligo). **(C-F)** Relative amounts of U-^13^C_6_-glucose-derived metabolites in glycolysis (C), the first round of TCA cycle (D), the PPP(E), and nucleotide synthesis (F) in DMSO-(black) or oligomycin-treated (orange) HSCs; In (B)-(F), biological replicates of the DMSO and oligomycin groups obtained on four separate days were pooled, analyzed by IC-MS, and quantified based on calibration curve data for each metabolite (see “**Ion chromatography mass spectrometry (IC-MS) analysis**” section in “**Methods**” for details). (G-H) Mito Stress test on the Seahorse flux analyzer for HSC and MyPs; ECAR (G) and OCR (H) before and after oligomycin treatment. (Data were obtained from n = 3 technical replicates for HSCs and n = 10 technical replicates for MyPs.) Data are shown as mean ± SD. * p ≤ 0.05, ** p ≤ 0.01, *** p ≤ 0.001 as determined by paired-samples t-test (C-E and G–H). Abbreviations: G6P, glucose-6-phosphate; F6P, fructose-6-phosphate; F1,6BP, fructose-1,6-bisphosphate; G3P, glycerol-3-phosphate; DHAP, dihydroxyacetone phosphate; 3PG, 3-phosphoglycerate; 2PG, 2-phosphoglycerate; PEP, phosphoenolpyruvate; PYR, pyruvate; LAC, lactate; Ac-CoA; acetyl-CoA; CIT, citrate; ACO, cis-aconitic acid, isocitrate; 2OG, 2-oxoglutarate; SUC, succinate; FUM, fumarate; MAL, malate; OAA, oxaloacetate; 6PG, glucose-6-phosphate; Ru5P, ribulose-5-phosphate; Xu5P, xylulose-5-phosphate; R5P, ribose-5-phosphate; S7P, sedoheptulose-7-phosphate; E4P, erythrose-4-phosphate; PRPP, phosphoribosyl pyrophosphate; IMP, inosine monophosphate; ATP, adenosine triphosphate; GTP, guanine triphosphate; UMP, uridine monophosphate; UTP, uridine triphosphate; TTP, thymidine triphosphate. See also Fig. S1–3.

### Phosphofructokinase (PFK) metabolism in HSCs is activated during proliferation and OXPHOS inhibition

To investigate whether glycolytic activation in HSCs after 5-FU treatment and OXPHOS inhibition could be demonstrated through unbiased mathematical simulations, we performed quantitative ^13^C metabolic flux analysis (^13^C-MFA). After generating a metabolic model for isotope labeling enrichment and setting appropriate lactate efflux values, a simulation was conducted using the labeled metabolite abundance data obtained from isotope tracer analysis. The appropriate lactate efflux for quiescent HSC (PBS-treated HSC) was determined to 65 after experimenting with values from 0–100. The lactate efflux of 5-FU- or oligomycin-treated HSCs was higher than that of quiescent HSCs based on the observation that labeled glycolytic metabolite levels were particularly elevated in *in vitro* tracer analysis (see “**Quantitative ^13^C-MFA with OpenMebius**” under "**Methods**" for more information). As a result, the variation in the flux values of all enzymatic reactions calculated in HSCs after 5-FU or oligomycin treatment became smaller compared to quiescent HSCs, suggesting that HSCs strictly regulated their metabolism in response to stress (Supplemental Figure 4A-C). Unlike PBS-treated HSCs, those treated with 5-FU or oligomycin exhibited preferential glycolytic activation rather than TCA- or PPP-based metabolic strategies; the first half of the glycolytic system appeared to be the site of metabolic activation (Figure 3A-J; Supplemental Figure 4D-U, Table S4). This increase in metabolic flux upstream of the glycolytic pathway was also supported by our *in vitr*o tracer analysis (Figure 1B and Figure 2B), suggesting that ^13^C-MFA was a valid metabolic simulation. Among the reactions in the first half of glycolysis, phosphorylation of fructose 6-phosphate by PFK is the irreversible and rate-limiting reaction^55^. A detailed review of *in vitro* isotope tracer analysis results showed that the ratio of fructose 1,6-bisphosphate (F1,6BP; the product of PFK) to F6P (the substrate of PFK) was greatly elevated in HSCs during proliferation and OXPHOS inhibition (Figure 3K-L). Together with the results of quantitative ^13^C-MFA, these findings suggested that HSCs exhibit elevated glycolytic flux relative to mitochondrial activity by increasing PFK enzyme activity under various stress conditions.

**Figure 3.**
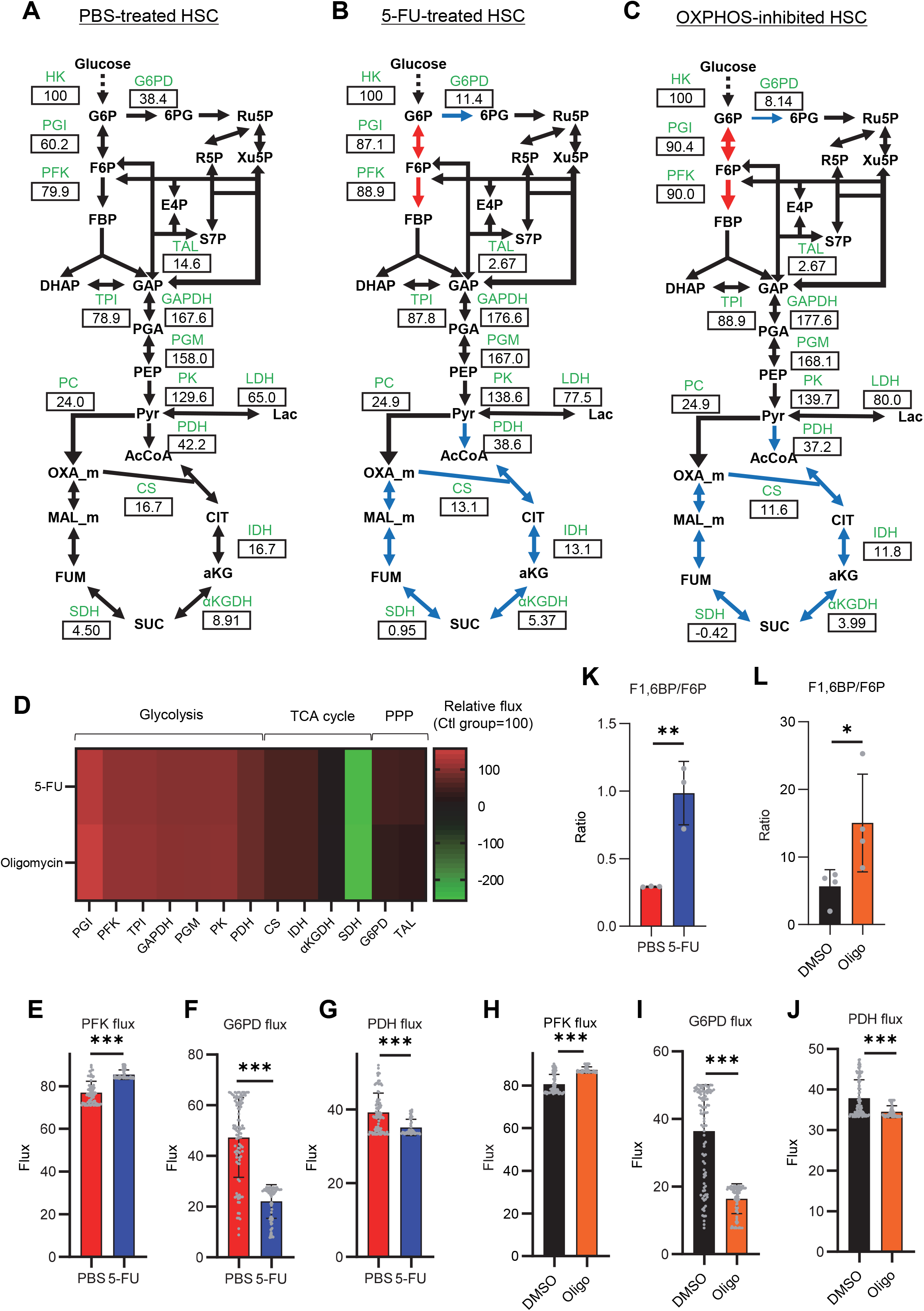
Quantitative ^13^C-MFA of quiescent, proliferative, and stressed HSCs. **(A-C)** Overview of quantitative ^13^C-MFA of PBS-treated HSCs (A), 5-FU-treated HSCs (B), and OXPHOS-inhibited HSCs (C). The representative net flux for each reaction with glucose uptake as 100 is shown in the squares below the catalytic enzymes for each reaction listed in green letters. Red arrows indicate reactions with particularly elevated fluxes and blue arrows indicate reactions with particularly decreased fluxes. (D) Heatmap of the relative flux of each enzyme in the 5-FU or oligomycin groups compared to that in the quiescent (Ctl) HSC (The metabolic flux of each enzyme in the Ctl group was standardized as 100.). **(E-J)** Fluxes due to reactions with PFK (E, H), G6PD (F, I), and PDH (G, J). Fluxes of HSCs derived from mice treated with 5-FU (blue bars) or PBS (red bars) (D-F) and of HSCs treated with DMSO (black bars) or oligomycin (orange bars) (G-I) are shown. Data is obtained from 100 simulations in OpenMebius, and flux data for each enzyme is displayed (Table S4). **(K-L)** Ratio of fructose 1,6-bisphosphate (F1,6BP) to fructose-6-phosphate (F6P) calculated from tracer experiments shown in Figure 1B and Figure 2B. Effects of 5-FU administration (K) or mitochondrial inhibition by oligomycin (L) are summarized. Data are shown as mean ± SD. * p ≤ 0.05, ** p ≤ 0.01, *** p ≤ 0.001 as determined by Student’s *t*-test (E-L). Abbreviations: HK, hexokinase; PGI, glucose-6-phosphate isomerase; PFK, phosphofructokinase; TPI, triose phosphate isomerase; GAPDH, glyceraldehyde-3-phosphate dehydrogenase; PGM, phosphoglycerate mutase; PK, pyruvate kinase; LDH, lactate dehydrogenase; PC, pyruvate carboxylase; PDH, pyruvate dehydrogenase; CS; citrate synthase; IDH, isocitrate dehydrogenase; αKGDH, α-ketoglutaric acid dehydrogenase; SDH, succinate dehydrogenase; G6PD, glucose-6-phosphate dehydrogenase; TAL, transaldolase. See also Fig. S4.

### HSCs under stress exhibit activation of glycolysis-initiated TCA cycle and NAS

To investigate the long-term glucose utilization of HSCs, we performed an *in vivo* tracer analysis with U-^13^C_6_ glucose based on recent reports^47,50^ (Supplemental Figure 5A; see “**Preparation and storage of *in vivo* U-^13^C_6_-glucose tracer samples**” under "**Methods**" for more information). In HSCs from 5-FU-treated mice, we observed increased labeling of glycolytic metabolites such as dihydroxyacetone phosphate, glycerol-3-phosphate, and phosphoenolpyruvate, as well as NAS metabolites such as inosine monophosphate and ATP, and those derived from TCA cycle such as aspartic acid and glutamate, compared to HSCs from PBS-treated mice (Supplemental Figure 5B-I, Table S5). When the amount of U-^13^C_6_-glucose-derived labeled metabolites in each pathway was calculated, more glucose-derived metabolites entered TCA cycle in the 5-FU-treated group than PBS-treated group (Supplemental Figure 5J). Thus, although short-term (10–30 min) *in vitro* tracer analysis showed that HSCs exhibited more potent activation of anaerobic glycolysis than of other pathways in response to 5-FU administration, long-term (approx. 3 h) labeling by *in vivo* tracer analysis revealed that glycolysis-initiated TCA cycle and NAS flux were activated in addition to enhanced anaerobic glycolysis. Importantly, despite differences in labeling times and supplementation of U-^13^C_6_ glucose metabolites from non-HSCs to HSCs *in vivo*, the activation of the glycolytic system was a common finding.

### PFKFB3 accelerates glycolytic ATP production during HSC cell cycling

*In vitro* and *in vivo* tracer analysis results collectively suggested that the activation of glycolysis catalyzed by PFK may have been the starting point for the activation of the entire HSC metabolism. To analyze the contribution of PFK to ATP metabolism in steady-state or stressed HSCs, we needed to develop an experimental system that could measure the dynamics of ATP concentrations in HSCs in a non-destructive, real-time manner. To this end, we used knock-in GO-ATeam2 mice as a FRET-based biosensor of ATP concentration (see “**Conversion of GO-ATeam2 fluorescence to ATP concentration**” under "**Methods**" for more information.). The number of bone marrow mononuclear cells (BMMNCs), as well as the frequency of HSCs (CD150^+^CD48^-^LSK) and other progenitor cells, in the bone marrow (BM) of GO-ATeam2^+^ mice were almost unchanged compared to of C57B6/J mice, except for a mild decrease in the Lin^-^ fraction (Supplementary Figure 6A-C). Using BMMNCs derived from GO-ATeam2^+^ mice, we developed a method to detect changes in ATP concentration with high temporal resolution when the activity of PFK was modulated (Supplemental Figure 6D-F). To validate our methods, we measured ATP concentrations in HSCs and MyPs with or without various nutrients (see “**Time-course analysis of FRET values**” under "**Methods**" for more information.). MyPs showed more rapid decreases in ATP concentration than HSCs, suggesting higher ATP consumption by progenitors (Supplemental Figure 6G-H). Adding glucose to the medium suppressed this decrease in MyPs; however, other metabolites (e.g., pyruvate, lactate, and fatty acids) had minimal effects, suggesting that ATP levels are glycolysis-dependent in MyPs (Supplemental Figure 6G-H), consistent with previous reports that the aerobic glycolytic enzyme M2 pyruvate kinase isoform (PKM2) is required for progenitor cell function^33^.

Further, we analyzed ATP consumption and metabolic dependency of cell-cycling HSCs after 5-FU administration (Figure 4A). After inhibiting glycolysis using 2-deoxy-D-glucose (2-DG) with other mitochondrial substrates, 5-FU-treated HSCs showed more rapid decreases in ATP concentration than PBS-treated HSCs (Figure 4B-C). In contrast, OXPHOS inhibition by oligomycin without glucose or mitochondrial substrates decreased the ATP concentration to a similar extent in both 5-FU- and PBS-treated HSCs, although 5-FU-treated HSCs showed earlier ATP exhaustion (Figure 4D-E). These data suggest that 5-FU-treated-HSCs upregulated ATP production via glycolysis, rather than relying on mitochondria. Apoptosis assay revealed a slight increase in early apoptotic cells (annexin V^+^ propidium iodide [PI]^-^) after 2-DG treatment and a slight decrease in the number of viable cells (Annexin V^-^ PI^-^) after oligomycin treatment, both to a very limited extent (approx. 5%) compared to the degree of ATP decrease, suggesting that the decrease in ATP after 2-DG or oligomycin treatment did not simply reflect cell death (Supplementary Figure 6I). Importantly, no metabolic changes in glycolysis or OXPHOS were observed in HSCs without cell cycle progression after 5-FU administration (very early phase: day 3; late phase: day 15) (Supplementary Figure 7A-H).

**Figure 4.**
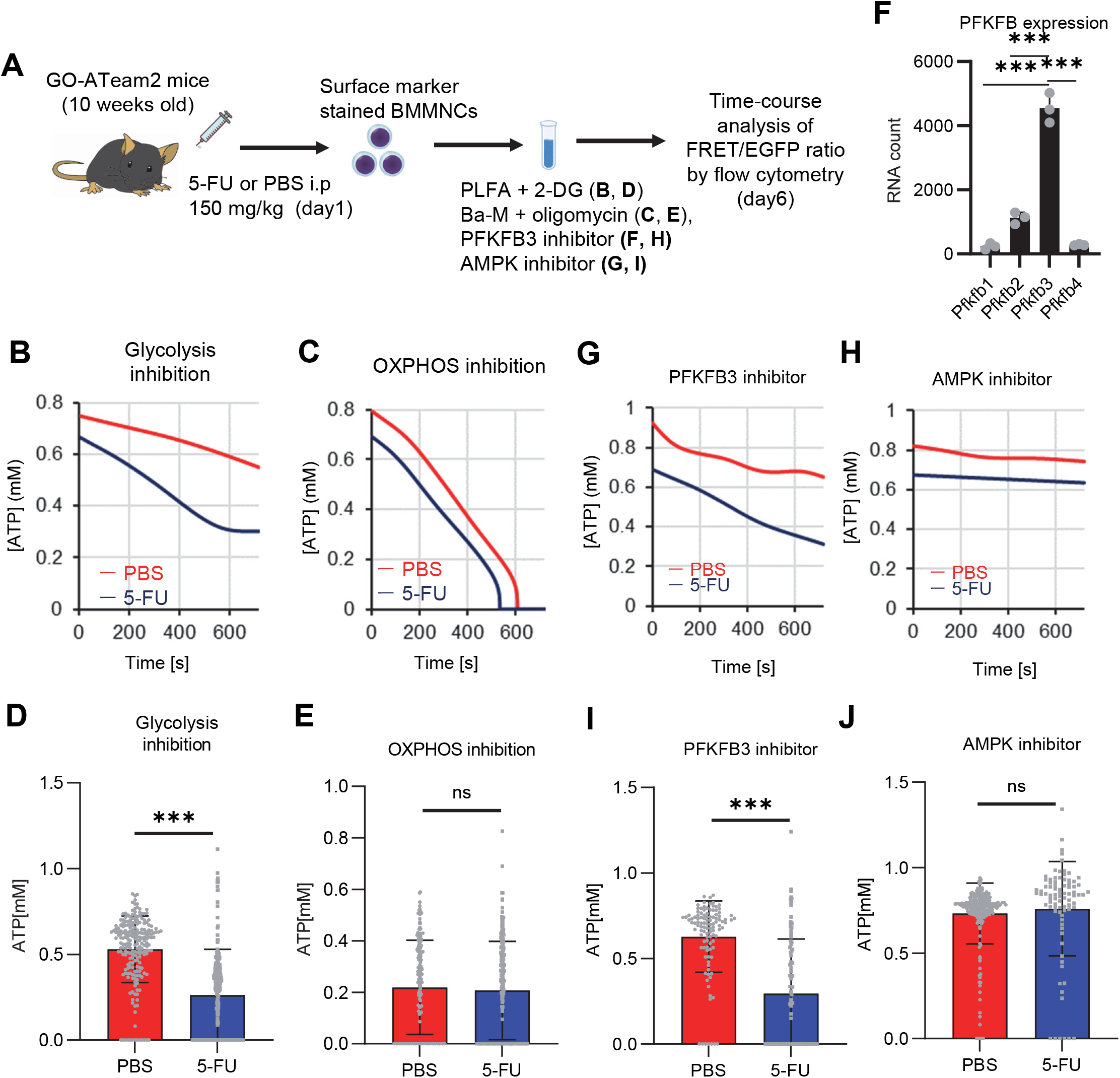
Pfkfb3 activates the glycolytic system in proliferating HSCs. **(A)** Experimental design used to conduct real-time ATP analysis of HSCs treated with 5-FU or PBS. PLFA medium containing mitochondrial substrates (pyruvate, lactate, fatty acids, and amino acids) but no glucose, was used for experiments with 2-DG; Ba-M containing neither mitochondrial substrates nor glucose was used for experiments with oligomycin, Pfkfb3 inhibitor, or AMPK inhibitor. **(B-I)** Results of real-time ATP analysis of PBS-(red) or 5-FU-treated (blue) HSCs after treatment with 2-DG (B, D), oligomycin (C, E), PFKFB3 inhibitor (F, H), or AMPK inhibitor (G, I). Bar graphs show corrected ATP concentrations for the last 2 min (D) of (B), 6–7 min (E) of (C), or the last 1 min (H, I) of (F, G) for PFKFB3 and AMPK inhibitors, respectively. Each group represents at least 60 cells. Data are representative results of pooled samples from three biological replicates. (see “**Time-course analysis of FRET values**” in “**Methods**” for details of the correction method used to calculate ATP concentration.) (J) Normalized mRNA counts of PFKFB isozymes based on the RNA sequencing of HSCs. Data are presented as mean ± SD. * p ≤ 0.05, ** p ≤ 0.01, *** p ≤ 0.001 as determined by Student’s *t*-test (D, E, H, and I) or a one-way ANOVA followed by Tukey’s test (J). See also Fig. S6–7.

PFK is allosterically activated by 6-phosphofructo-2-kinase/fructose-2,6-bisphosphatase (PFKFB). Among the four isozymes of mammalian PFKFB, PFKFB3 is the most favorable for PFK activation^56^, and is the most highly expressed in HSCs (Figure 4F). Therefore, we investigated whether PFKFB3 contributes to glycolytic plasticity in HSCs during proliferation. When treated with the PFKFB3-specific inhibitor AZ PFKFB3 26^57^, compared with HSCs from PBS-treated mice, HSCs from 5-FU-treated mice showed decreased ATP levels (Figure 4G, I; Supplementary Figure 7I). Although AMPK activates PFKFB3 in other contexts^58^, AMPK inhibition by dorsomorphin did not alter ATP concentration in 5-FU-treated-HSCs (Figure 4H, J).

Finally, we investigated the nutrients that drive OXPHOS in PBS- or 5-FU-treated HSCs. Exposure of PBS- or 5-FU-treated HSCs to either etomoxir, a FAO inhibitor, or 6-diazo-5-oxo-L-norleucine (DON), a glutaminolysis inhibitor, alone or in combination, did not decrease ATP concentrations (Supplementary Figure 7J-M). Subsequent assessment of FAO activity using FAOBlue, a fluorescent probe for the FAO activity assay^59^, showed no significant differences between PBS- and 5-FU-treated HSCs (Supplementary Figure 7N). Thus, neither FAO nor glutaminolysis appeared to be essential for the short-term maintenance of ATP levels in cell-cycling HSCs after 5-FU administration. Notably, the addition of glucose and a Pfkfb3 inhibitor to etomoxir rapidly reduced ATP concentrations in HSCs (Supplementary Figure 7O-P). This suggests that etomoxir may partially mimic the effects of oligomycin, indicating that OXPHOS is primarily driven by FAO, but can be compensated by Pfkfb3-accelerated glycolysis in HSCs. Conversely, exposure of HSCs to DON in combination with a Pfkfb3 inhibitor did not decrease ATP concentrations (Supplementary Figure 7O-P), suggesting that ATP production via glutaminolysis is limited in HSCs.

### OXPHOS inhibition accelerates glycolysis to sustain ATP levels in HSCs, but not in progenitors

To assess differences in metabolic dependence between steady-state or stressed and naturally proliferating HPCs, we altered ATP metabolism in HSCs and progenitors using 2-DG or oligomycin (Figure 5A). Oligomycin treatment rapidly depleted ATP in HSCs and all HPC fractions (green lines in Figure 5B-C; Supplemental Figure 8A-D). Treatment with 2-DG decreased ATP concentrations for a short amount of time (∼12 min) in HSCs and HPCs, but ATP reduction was less evident than that induced by oligomycin (blue lines in Figure 5B-C; Supplemental Figure 8A-D). The ATP reduction induced by 2-DG treatment was particularly low (∼15%) in HSCs, multipotent progenitor cells (MPPs), and common lymphoid progenitors (CLPs) relative to that in common myeloid progenitors (CMPs), granulocytes-macrophage progenitors (GMPs), and megakaryocyte-erythrocyte progenitors (MEPs) (Figure 5D).

**Figure 5.**
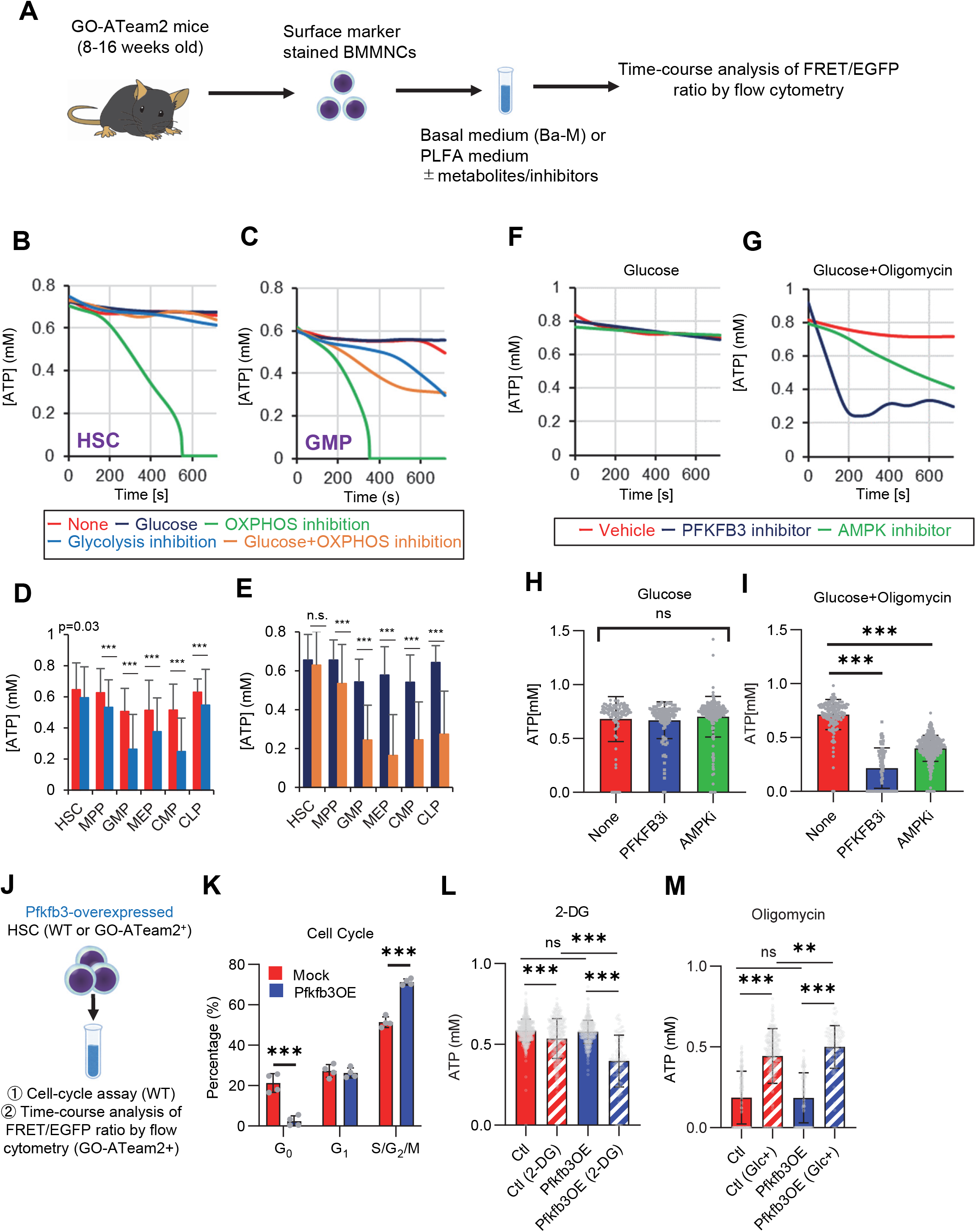
PFKFB3 accelerates glycolysis in HSCs under OXPHOS inhibition in an AMPK-dependent manner. **(A)** Experimental design of real-time ATP analysis using GO-ATeam2 knock-in BMMNCs. Ba-M was used in experiments with oligomycin. For other experiments, PLFA medium was used. **(B-C)** Evaluation of factors affecting ATP concentration in HSCs (B) and GMPs (C) based on the GO-ATeam2 system. GO-ATeam2 knock-in BMMNCs were incubated with glucose, oligomycin, 2-DG, or glucose plus oligomycin, and the FRET/EGFP ratio was calculated. **(D)** ATP concentration in indicated stem/progenitor fractions in PLFA medium (red bars) alone or PLFA medium plus 2-DG (blue bars). ATP concentration for the last 2 min of the analysis time is shown. Data is summarized from Figure 5B-C and Supplemental Figure 5A-D. Each group represents at least 110 cells. Data are representative results of pooled samples from three biological replicates. **(E)** ATP concentration in indicated stem/progenitor fractions in Ba-M plus glucose (dark blue bars) or Ba-M plus glucose and oligomycin (orange bars). ATP concentration for the last 1 min of the analysis period is shown. Data is summarized from Figure 5B-C and Supplemental Figure 5A-D. Each group represents at least 43 cells. Data are representative results of pooled samples from three biological replicates. **(F-I)** Effects of PFKFB3 or AMPK inhibitors (PFKFB3i or AMPKi, respectively) on ATP concentration in HSCs from GO-ATeam2 mice in Ba-M plus glucose only (F) or Ba-M plus glucose and oligomycin (G). ATP concentrations for the last 1 min of the analysis period are shown in (H) and (I) for glucose only and glucose with oligomycin groups, respectively. Each group represents at least 90 cells. Data are representative results of pooled samples from three biological replicates. **(J)** Experimental schema for cell cycle assay and real-time ATP concentration analysis after overexpression of *Pfkfb3*. **(K)** Cell cycle status of *Pfkfb3*-overexpressing (*Pfkfb3*OE) and *mock*-transduced HSCs. **(L-M)** Effects of inhibitors on ATP concentration in *Pfkfb3*-overexpressing GO-ATeam2^+^ HSCs. Cells were exposed to vehicle or 2-DG (L), oligomycin in the presence or absence of glucose 12.5 mg/dL (M), and ATP concentrations for the last 2 min (L) or 1 min (M) of the analysis period were calculated. Data are representative results of pooled samples from three biological replicates. Data are presented as mean ± SD. * p ≤ 0.05, ** p ≤ 0.01, *** p ≤ 0.001 as determined by Student’s *t*-test (D, E, and K) or one-way ANOVA followed by Tukey’s test (H, I, L, and M). See also Fig. S8.

Next, we investigated the role of glycolysis in ATP production during OXPHOS inhibition by combining oligomycin administration and glucose supplementation. ATP concentration remained more stable in HSCs treated with oligomycin and glucose than in those treated only with oligomycin. Similar results were not seen in HPCs, indicating that HSCs have the plasticity to upregulate glycolytic ATP production to meet demands (orange lines in Figure 5B-C; Supplemental Figure 8A-D, summarized in Figure 5E). Similar to oligomycin treatment, rotenone (Complex I inhibitor) and carbonyl cyanide 4-(trifluoromethoxy)phenylhydrazone (FCCP, mitochondrial uncoupler) treatments, which inhibit OXPHOS-derived ATP production, also decreased ATP concentrations in HSCs, but not when administered simultaneously with glucose (Supplementary Figure 8E-F). Furthermore, with oligomycin, HSCs, but not HPCs, maintained ATP concentrations at low glucose levels (50 mg/dL) (Supplemental Figure 8G). These analyses suggest that ATP was produced by mitochondrial OXPHOS in steady-state HSCs, and that only HSCs, but not HPCs, maintained ATP production by glycolysis when OXPHOS was compromised.

### PFKFB3 accelerates glycolytic ATP production during OXPHOS inhibition

Next, to understand whether PFKFB3 contributes to ATP production in HSCs under OXPHOS inhibition, we evaluated PFKFB3 function under OXPHOS inhibition using the GO-ATeam2^+^ BMMNCs. In oligomycin-treated HSCs, PFKFB3 inhibition led to rapidly decreased ATP concentration that was not observed in HSCs not treated with oligomycin (Figure 5F-I). We examined the effects of HSPC metabolic regulators on ATP levels in oligomycin-treated HSCs. Inhibiting PKM2, which accelerates glycolysis in steady-state progenitors^33^, significantly reduced ATP levels in oligomycin-treated HSCs (Supplemental Figure 8H, J). Inhibiting LKB1, a kinase upstream of AMPK^60,61^, did not affect the ATP concentration in oligomycin-treated HSCs (Supplemental Figure 8I, K), whereas levels of adenosine monophosphate (AMP), which also activates AMPK, increased in oligomycin-treated but not in 5-FU-treated HSCs (Supplemental Figure 8L). This may explain differences in AMPK-dependent ATP production between proliferative HSCs and HSCs under OXPHOS inhibition.

Next, we tested the effects of PFKFB3 on ATP concentration in HPCs. Unlike HSCs, HPCs exhibited PFKFB3-dependent ATP production, even without oligomycin (Supplemental Figure 8M-Q). Therefore, ATP production in steady-state HSCs was PFKFB3-independent, and proliferative stimulation or OXPHOS inhibition plastically activated glycolytic ATP production in a PFKFB3-dependent manner to meet ATP demand.

### PFKFB3 activity renders HSCs dependent on glycolysis

Next, we investigated whether PFKFB3 activity itself confers glycolytic dependence on HSCs. We retrovirally overexpressed *Pfkfb3* in HSCs and performed cell cycle analysis (Figure 5J). *Pfkfb3*-overexpressed HSCs increased the proportion of cells in the S/G2/M phase and decreased the number of G_0_ cells compared to *mock*-overexpressed HSCs (Figure 5K). Next, we retrovirally overexpressed *Pfkfb3* in GO-ATeam2^+^ HSCs and performed real-time ATP measurement (Figure 5J). *Pfkfb3*-overexpressing GO-ATeam2^+^ HSCs did not show changes in ATP concentrations relative to those in *mock*-transduced cells (Figure 5L; Supplemental Figure 8R). Upon 2-DG treatment, *Pfkfb3*-overexpressing HSCs showed a greater decrease in ATP concentration than *mock*-transduced HSCs did (Figure 5L; Supplemental Figure 8S). However, oligomycin treatment of both *mock*-transduced and *Pfkfb3*-overexpressing HSCs decreased ATP concentration to comparable levels (Figure 5M; Supplemental Figure 8T). Notably, *Pfkfb3*-overexpressing HSCs recovered ATP levels more effectively under low glucose conditions (12.5 mg/dL) than did *mock*-transduced HSCs (Figure 5M; Supplemental Figure 8U). These data suggest that PFKFB3 directly conferred glycolytic dependence onto HSCs by modulating the cell cycle and increasing their ATP-generating capacity via glycolysis under metabolic stress.

### PFKFB3 methylation by PRMT1 supports ATP production by cell-cycling HSCs

Next, we investigated how 5-FU-treated-HSCs regulate PFKFB3 independently of AMPK (Figure 4G-J). PFKFB3 activity is regulated at multiple levels^62^, and PFKFB3 transcript and protein levels in HSCs remained unchanged during 5-FU-induced cell cycling (Figure 6A-B). Phosphorylation can also regulate PFKFB3 activity^58,63,64^; however, we observed no change in PFKFB3 phosphorylation in 5-FU-treated-HSCs (Figure 6C). Upon oligomycin exposure, PFKFB3 was phosphorylated by AMPK in the HSCs (Figure 6D). PFKFB3 is also methylated, and its activity is upregulated by protein arginine methyltransferase 1 (PRMT1)^65^. We observed that *Prmt1* expression increased in 5-FU-treated-HSCs relative to that in PBS-treated-HSCs (Figure 6E). Furthermore, PFKFB3 methylation was significantly induced in 5-FU-treated-HSCs than in PBS-treated-HSCs (Figure 6F). Treatment of HSCs with a PRMT1 inhibitor decreased PFKFB3 methylation (Figure 6G), suggesting that PRMT1 catalyzed PFKFB3 methylation. In contrast, the number of transcripts regulated by PRMT1 decreased or was unchanged (Supplementary Figure 9), suggesting that the transcriptional regulatory function of PRMT1 is limited. To investigate whether glycolytic activity in HSCs was regulated by m-PFKFB3, mice treated with PBS or 5-FU were injected with 2-NBDG, and m-PFKFB3 levels in HSCs with high and low 2-NBDG uptake were quantified. Regardless of PBS or 5-FU treatment, HSCs with high 2-NBDG uptake exhibited higher m-PFKFB3 levels than those with low uptake (Figure 6H), suggesting that m-PFKFB3 regulated the activity of the glycolytic system in HSCs.

**Figure 6.**
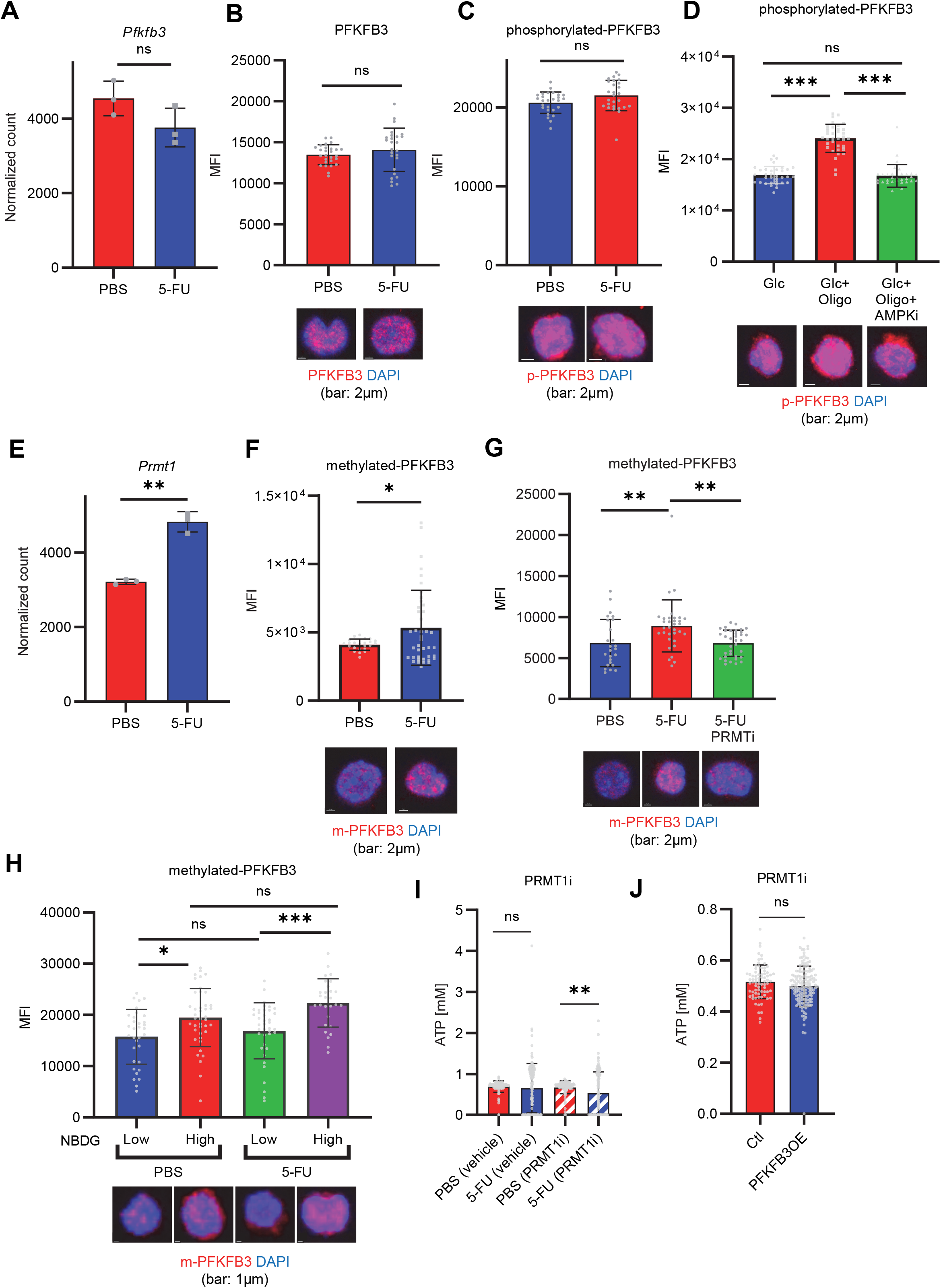
PFKFB3 methylation by PRMT1 enables ATP production by cell-cycling HSCs. **(A)** Normalized *Pfkfb3* mRNA counts based on RNA sequencing of PBS-treated (red) or 5-FU-treated (blue) HSCs. Data are representative results of pooled samples from three biological replicates. Data were extracted from the same pooled samples as in Figure 4J and Figure S9. **(B)** Quantification of mean fluorescent intensity (MFI) of PFKFB3 protein in PBS- or 5-FU-treated HSCs. The lower part of the graph shows representative images of immunocytochemistry of PFKFB3 in each group. n = 26–27 single HSCs for each group. The data are representative results from two independent experiments. **(C)** Quantification of MFI of phosphorylated-PFKFB3 (p-PFKFB3) protein in PBS- or 5-FU-treated HSCs. The lower part of the graph shows representative images of immunocytochemistry of p-PFKFB3 in each group. n = 27 single HSCs for each group. The data are representative results from two independent experiments. **(D)** Quantification of mean fluorescence intensity (MFI) of p-PFKFB3 in HSCs treated with glucose (200mg/dL); glucose plus oligomycin (1 µM); and glucose, oligomycin, and dorsomorphin (100 µM) for 5 min. The lower part of the graph shows representative images of immunocytochemistry of p-PFKFB3 in each group. n = 32–36 for each group. The data are representative results from two independent experiments. **(E)** Normalized *Prmt1* mRNA counts based on RNA sequencing of PBS-treated (red) or 5-FU-treated (blue) HSCs. Data are representative results of pooled samples from three biological replicates. **(F)** MFI quantification of methylated-PFKFB3 (m-PFKFB3) in PBS- or 5-FU-treated HSCs. The lower part of the graph shows representative images of immunocytochemistry of m-PFKFB3 in each group. n = 23–41 for each group. The data are representative results from three independent experiments. **(G)** Quantification of MFI of m-PFKFB3 in PBS- or 5-FU-treated HSCs or 5-FU-treated HSCs after 15 min treatment with a PRMT1 inhibitor (90 μg/mL GSK3368715); n = 25–35 single HSCs for each group. The lower part of the graph shows representative images showing immunocytochemistry of m-PFKFB3. Data represent a single experiment. **(H)** Quantitation of m-PFKFB3 in NBDG-positive or -negative HSCs in mice treated with PBS or 5-FU. The lower part of the graph shows representative images of immunocytochemistry of m-PFKFB3 in each group. n = 28–41 for each group. The data are representative results from two independent experiments. **(I)** Corrected ATP levels in PBS-(red) or 5-FU-treated (blue) HSCs 15 min after treatment with vehicle or a PRMT1 inhibitor (90 µg/mL GSK3368715). Each group represents at least 101 cells. Data are representative results of pooled samples of two biological replicates. (see “**Time-course analysis of FRET values**” in “**Methods**” for details of the correction method used to calculate ATP concentration.) (**J**) ATP concentration in mock-transduced (Ctl) or PFKFB3-overexpressed (OE) HSCs after treatment with the PRMT1 inhibitor (90 µg/mL GSK3368715). ATP concentration for the last 1 min of the analysis period is shown. Data are presented as mean ± SD. * p ≤ 0.05, ** p ≤ 0.01, *** p ≤ 0.001 as determined by Student’s *t*-test (A-C, E-F, and I-J) or one-way ANOVA followed by Tukey’s test (D, G, and H). See also Fig. S9.

Further, we analyzed the potential effects of PRMT1 inhibition on ATP concentration in GO-ATeam2^+^ HSCs. Treatment with the PRMT1 inhibitor significantly decreased ATP levels in 5-FU-treated-HSCs than in PBS-treated-HSCs (Figure 6I). In contrast, the retroviral overexpression of *Pfkfb3* in GO-ATeam2^+^ HSCs abolished the effect of the PRMT1 inhibitor on ATP reduction (Figure 6J). These findings indicated that ATP levels in 5-FU-treated-HSCs were supported by PRMT1 methylation–mediated PFKFB3 activation.

### PFKFB3 contributes to HSPC pool expansion and stress hematopoiesis maintenance

Finally, we analyzed PFKFB3 function in HSCs during hematopoiesis. We cultured HSCs with a PFKFB3 inhibitor *in vitro* under quiescence-maintaining or proliferative conditions (Supplementary Figure 10A)^54^. Cell count in HSC-derived colonies decreased following treatment with a PFKFB3 inhibitor under proliferative, but not quiescence-maintaining, conditions (Supplementary Figure 10B). We also knocked out *Pfkfb3* in HSCs using the less toxic, vector-free CRISPR-Cas9 system and cultured the cells under quiescence-maintaining or proliferative conditions (Supplementary Figure 10A) based on recent reports by Shiroshita et al.^66^. Again, cell numbers in *Pfkfb3*-knockout (KO) HSC–derived colonies decreased only in proliferative cultures when compared to control cultures (*Rosa26*-KO HSCs) (Supplementary Figure 10C, E, F). We retrovirally overexpressed *Pfkfb3* in HSCs and cultured them under quiescence maintenance or proliferative conditions (Supplementary Figure 10A). *Pfkfb3*-overexpressing HSC colonies showed increased cell count compared to that of *mock*-transduced cells, but only under proliferative conditions (Supplementary Figure 10D).

To assess PFKFB3 function in HSCs *in vivo*, we transplanted *Pfkfb3*-KO HSCs (Ly5.2^+^) or wild type (WT) control HSCs into lethally irradiated recipients (Ly5.1^+^) as well as Ly5.1^+^ competitor cells (Figure 7A), and the behavior of *Pfkfb3*-KO cells was evaluated by Sanger sequencing of peripheral blood (PB) cells^66^. In the KO group, donor-derived chimerism in PB cells decreased relative to that in the WT control group during the early phase (1 month post-transplant) but recovered thereafter (Figure 7B). Next, we retrovirally transduced Ly5.2^+^ HSCs with *Pfkfb3* S461E (*Pfkfb3*CA), a constitutively active PFKFB3 mutant, and transplanted them into lethally irradiated recipients (Ly5.2^+^), along with Ly5.1^+^ competitor cells (Figure 7A, Supplementary Figure 10G). Donor chimerism during the early post-transplant period in the *Pfkfb3*CA-overexpressing group was significantly higher than that in the *mock*-transduced group (Figure 7C). These findings suggest that PFKFB3 may play a role in the differentiation and proliferation of HSCs. Therefore, we compared the contribution of PFKFB3 to HSPC function at steady state and after myeloproliferative stimulation. *Pfkfb3*- or *Rosa*-KO HSPCs were transplanted into recipients (Ly5.1^+^). After 2 months, recipients received 5-FU intraperitoneally, and the dynamics of *Pfkfb3*- or *Rosa*-KO cell abundance in PB was assessed (Figure 7D). In PB cells prior to 5-FU administration, *Pfkfb3*- or *Rosa*-KO HSPC-derived blood cells were almost equally present, suggesting a limited involvement of PFKFB3 in steady-state blood cell production (Figure 7E). However, after 5-FU administration, *Pfkfb3*-KO HSPC-derived blood cell abundance was reduced compared to that in the *Rosa* group (Figure 7E). This change occurred on day 6 after 5-FU administration (day 1), when the cell cycle of HSCs was activated (Supplementary Figure 1D), supporting the idea that PFKFB3 contributes to HSC proliferation and differentiation into HSPCs.

**Figure 7.**
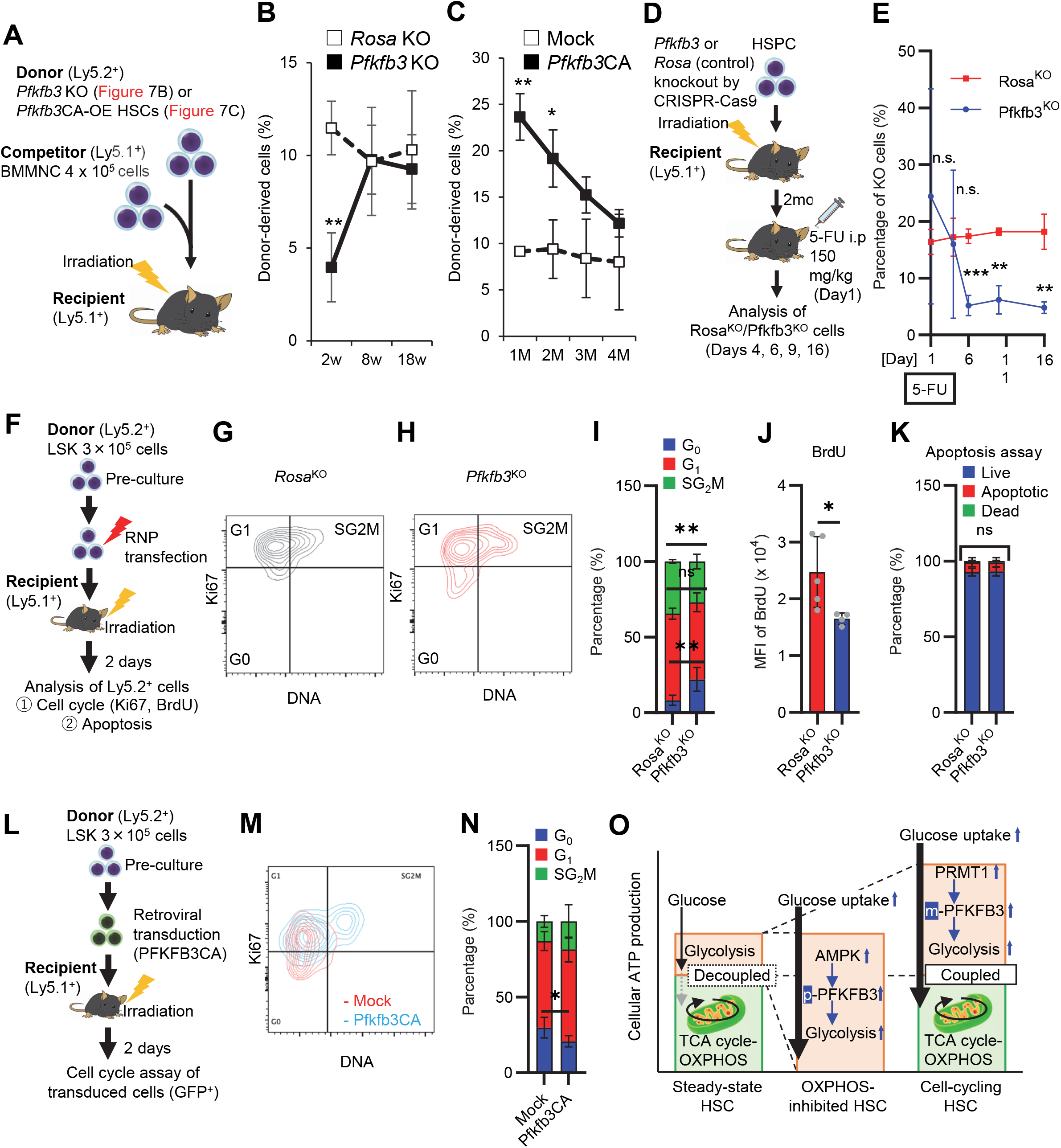
PFKFB3 maintains HSC function under proliferative stress. **(A-C)** Transplant analysis of *Pfkfb3-*KO or *Pfkfb3*CA-overexpressing HSCs. Experimental design (B). PB chimerism of donor-derived cells at 4 months post-transplant. *Pfkfb3-*KO group, n = 6; *Rosa-*KO group, n = 4; (B) *Pfkfb3* group, n = 5; pMY-IRES-GFP group, n = 4. (C) The data are representative results from two independent experiments. **(D-E)** 5-FU administration after bone marrow reconstruction with *Pfkfb3*- or *Rosa*-KO HSPCs. Experimental schema (D). Behavior of the *Pfkfb3*- or *Rosa*-KO cells in PB after 5-FU administration (E). n = 5 for each group. **(F-K)** Cell cycle analysis and apoptosis assay of *Pfkfb3*- or *Rosa*-KO HSPCs on day 2 post-BMT. Experimental schema (F). Representative plots of Ki67/Hoechst33432 staining of *Rosa*-KO (G) or *Pfkfb3*-KO (H) HSPCs and summary of analysis (I); summary of *in vivo* BrdU labeling assay (J). Apoptosis assay results (K). n = 4–5 biological replicates for each group. **(L-N)** Cell cycle analysis of *Pfkfb3*CA or *Mock*-overexpressing HSPCs on day 2 after BMT. Experimental Schema (L). Representative plot of Ki67/Hoechst33432 staining for both groups (M) and summary of analysis (N). n = 5 biological replicates for each group. **(O)** Models showing ATP production and regulation in quiescent, OXPHOS-inhibited, and cell-cycling HSCs. Note that the GO-ATeam2 system identified plastic acceleration of glycolysis by PFKFB3 in response to different types of stress maintains ATP levels. Data are presented as mean ± SD. * p ≤ 0.05, ** p ≤ 0.01, *** p ≤ 0.001 as determined by Student’s *t*-test (B, C, E, H, J, K, and N). See also Fig. S10.

To investigate the mechanisms underlying the short-term effects of PFKFB3 on hematopoiesis after bone marrow transplantation (BMT), we evaluated cell cycle and apoptosis of *Pfkfb3*-KO or -overexpressing HSPCs on day 2 after BMT (Figure 7F). Cell cycle was analyzed by Ki67/Hoechst33432 staining and *in vivo* BrdU labeling ^50^, which showed that cell cycle progression was suppressed in *Pfkfb3*-KO HSPCs (Figure 7G-J). In contrast, *Pfkfb3*-KO cells did not show increased apoptotic rates or decreased homing efficiency after BMT (Figure 7K; Supplementary Figure 10H). Furthermore, we examined the cell cycle of HSPCs overexpressing *Pfkfb3*CA on day 2 after BMT (Figure 7L) and found that *Pfkfb3*CA-overexpressing HSPCs showed accelerated cell cycle compared to *mock*-overexpressing HSPCs (Figure 7M-N). These data suggest that PFKFB3 contributes to HSC proliferation and differentiates cell production in *in vitro* and *in vivo* proliferative environments (cytokine stimulation and transplantation).

## Discussion

In this study, by combining metabolomic tracing of U-^13^C_6_-labeled glucose and ^13^C-MFA, we quantitatively identified the metabolic programs used by HSCs during steady-state, cell-cycling, and OXPHOS inhibition. Under proliferative stress, HSCs uniformly shift from mitochondrial respiration to glycolytic ATP production and PPP activation, which represent hallmarks of cell-cycling mammalian cells^67^. Previous reports have emphasized the importance of glycolysis in maintaining HSC quiescence, but have primarily analyzed HSCs in transplant assays, wherein HSCs must enter the cell cycle^32,68^. Prior analysis of repopulation capacity, which is positively correlated with enhanced glycolysis, may have overestimated glycolytic ATP production and overlooked mitochondrial ATP production during native hematopoiesis. In fact, some studies have suggested that OXPHOS activity is important for HSC maintenance and function^21^.

Our method was based on recently reported quantitative metabolic analysis techniques for very small numbers of cells^23,46–50^, such as HSCs, and expands our knowledge of HSC metabolism during stress hematopoiesis. In our study, 5-FU administration in mice transiently decreased ATP concentration in HSCs in parallel with cell cycle progression, suggesting that HSC differentiation and cell cycle progression are closely related to intracellular metabolism and can be monitored by measuring ATP concentration. We mainly analyzed a mixture of EPCR^+^ and EPCR^-^ HSCs, and we believe that the observed cell cycle progression and promotion of glycolysis in both EPCR^+^ and EPCR^-^ HSCs support the validity of our claims (Figure 1L, Supplementary Figure 1G-K). According to ^13^C-MFA enzymatic reaction flux of PFK in 5-FU-treated HSCs indicated a relative increase of approximately 10%. However, the flux value obtained by ^13^C-MFA was calculated with glucose uptake as 100. Thus, when combined with the overall increase in the glycolytic pool demonstrated by *in vitro* isotopic glucose tracer analysis and *in vivo* NBDG analysis, rapid acceleration of glycolysis becomes evident throughout the HSCs, including subpopulations that were less responsive to stress^43–45^. These findings are consistent with reports suggesting that HSCs have relatively low biosynthetic activity^69,70^ that is rapidly activated in response to cell proliferation stimuli^40,71^. Notably, we found that HSCs could accelerate glycolytic ATP production to fully compensate for mitochondrial ATP production under OXPHOS inhibition, a phenomenon that is difficult to identify without real-time ATP analysis. Thus, HSCs exposed to acute stresses choose to change the efficiency of glucose utilization (accelerated glycolytic ATP production) rather than other energy sources. *In vivo*, a completely glucose-deficient environment is improbable. Therefore, even under conditions such as hypoxia, where OXPHOS is inhibited, it is conceivable that glycolysis is accelerated to maintain ATP concentrations. Glucose tracer analysis showed NAS suppression under OXPHOS inhibition, leading to glycolysis without cell proliferation (Figure 2C-F; Supplementary Figure 1N). This suppression can be attributed to several factors: phosphates derived from ATP are added to nucleotide mono-/di-phosphates during NAS; the primary source of ATP production, OXPHOS, is impaired; and the presence of enzymes, such as dihydroorotate dehydrogenase, which are conjugated with OXPHOS ^72^. Such multifactorial effects raise new questions about the relationship between OXPHOS and nucleotide synthesis. On the other hand, we observed that ATP production in steady-state or cell-cycling HSCs and in naturally proliferating HPCs depended more on mitochondrial OXPHOS than on glycolysis; inhibiting glycolysis in steady-state HSCs resulted in only mild ATP decreases, suggesting that OXPHOS is still the major source of ATP production even in a medium saturated with hypoxia mimicking the BM environment. The p50 value of mitochondria (the partial pressure of oxygen at which respiration is half maximal) is less than 0.1 kPa, corresponding to an oxygen concentration of less than 0.1% under atmospheric pressure ^73^, suggesting that even under hypoxic conditions, OXPHOS can maintain some level of activity. Because FAO and the mitochondrial respiratory chain are necessary for HSC self-renewal and quiescence^21,37,39,54^, fatty acids may support mitochondrial ATP production independently of fluxes from glycolysis. FAO and glutaminolysis were not immediately essential for ATP production in HSCs. Given reports on the long-term necessity of FAO and glutaminolysis for HSC maintenance ^39,74^, ATP concentrations could be maintained in the short term by compensatory pathways. Furthermore, although glycolysis and TCA cycle are decoupled in steady-state HSCs, in response to cell cycle progression, anaerobic glycolytic metabolism in HSCs is enhanced (Figure 1) and fluxes to TCA cycle and PPP from the glycolytic system are also promoted (Supplementary Figure 5). The degree of glycolysis and TCA cycle coupling observed by *in vitro* and *in vivo* tracer analysis differed, likely due to differences in labeling time (10–30 min *in vitro* and 3 h *in vivo*). In particular, *in vivo* tracer analysis allows all cells to be capable of metabolizing U-^13^C_6_-glucose and providing its metabolites to HSCs, and there is a significant amount of time, approximately 120–180 minutes, after glucose labeling to purify HSCs. Metabolic reactions will continue during this time and subsequent processing on ice, which may increase the influx of labeled carbon into the TCA cycle. This complex dynamic in the *in vivo* tracer analysis makes it difficult to determine whether the labeled carbon influx is the result of direct influx from glycolysis or the re-uptake of metabolites by HSCs that have been processed by other cells. This is in contrast to *in vitro* analysis where such extended metabolic processing does not occur. Furthermore, despite an increased carbon influx into the TCA cycle *in vivo*, ATP production from mitochondria does not show a corresponding increase after 5-FU treatment, as shown by the GO-ATeam2 analysis shown in Figure 4C. Despite these technical differences, an essential common finding from both *in vivo* and *in vitro* analyses is the activation of glycolysis and nucleotide synthesis (NAS) in 5-FU-treated HSCs, highlighting critical metabolic changes in response to treatment. Moreover, these data provide direct evidence that glycolysis and TCA cycle become functionally uncoupled in quiescent HSCs^32,38^. Our findings are also consistent with previous reports of OXPHOS activation associated with HSC proliferation^32,36,75,76^. In other words, HSCs exhibit an increased proportion of anaerobic glycolysis–derived ATP by PFKFB3 upon proliferation and OXPHOS inhibition; furthermore, the glycolytic system is the starting point of metabolic activation and is indispensable for the overall enhancement of HSC metabolism (Figure 7H).

HPCs and leukemic cells accelerate glycolytic ATP production using PKM2 for differentiation and transformation, respectively^33^; however, we demonstrated that glycolytic acceleration does not fully compensate for mitochondrial ATP production in HPCs. Mechanistically, PFKFB3 increased glycolytic activity in HSCs to maintain ATP concentrations during proliferation and OXPHOS inhibition. Furthermore, inhibition of PFKFB3 in addition to OXPHOS does not result in a complete loss of ATP in HSCs, suggesting the robustness of HSC metabolism (Figure 5G). Under steady-state conditions, naturally proliferating HPCs rely on PFKFB3 for ATP production, whereas HSCs do not. This may explain the reduction of ECAR after oligomycin treatment in MyPs as shown by the Mito stress test (Figure 2G). In other words, while PFKFB3-dependent active glycolysis and mitochondria must always be coupled in MyPs, this is not necessarily the case in HSCs, even after 5-FU treatment (Figure 1G). Therefore, we can infer that quiescent HSCs at steady state can produce ATP via PFKFB3 activation in response to stress, enabling additional ATP generation. Furthermore, overexpression of *Pfkfb3* in HSCs increased glycolytic dependency, suggesting that PFKFB3 itself can modulate metabolic dependency in HSCs. Changes in glycolytic dependency in HSCs overexpressing *Pfkfb3* may seem small (0.06–0.13 mM) (Figure 5L, M). However, it is noteworthy that the rate of the reaction catalyzed by PFK varies greatly within a very narrow range of ATP concentrations, less than 1 mM. Webb et al. analyzed the factors controlling PFK activity and reported that the reaction rate of PFK varies by approximately 40% in the 0.3–1 mM ATP concentration range^77^. The reason that differences in glycolytic dependence could be detected in cells overexpressing *Pfkfb3* may be that the ATP concentration at the time of analysis was approximately 0.5–0.6 mM, which is within the range where a small change in ATP concentration can dynamically alter PFK activity.

PFKFB3 supports hematopoiesis in contexts that require robust HSPC proliferation *in vitro* and *in vivo*. We showed that the positive or negative effect of *Pfkfb3* overexpression or KO on differentiated blood cell production is gradually lost after BMT. This is because HSPCs require PFKFB3 for cell cycle progression during stress hematopoiesis in the early phase after BMT (Figure 7F-J, and L-N). However, even during stress hematopoiesis, PFKFB3 is not involved in cell death or homing efficiency (Figure 7K; Supplementary Figure 10H) and appears to contribute primarily to the regulation of transient HSPC proliferation in the BM cavity. HSCs no longer require PFKFB3 for a certain period of time after BMT, probably because they regain a quiescent state. This is consistent with the fact that inhibition of PFKFB3 in quiescent HSCs does not reduce the ATP concentration (Figure 5F, H), suggesting that the activity of PFKFB3 is plastically modified. HSC metabolic plasticity is also illustrated by the mode of PFKFB3 activation, differing depending on stress type. During proliferative stress, PRMT1 methylates PFKFB3 in the HSCs to promote glycolytic ATP production, a modification that increases its activity^65^. PRMT1 is required for stress hematopoiesis^78^, but its downstream targets in HSCs remain unclear. Our results strongly suggest that PRMT1 targets PFKFB3 to stimulate glycolysis in HSCs. In contrast, under OXPHOS inhibition, PFKFB3 phosphorylation by AMPK is induced—another modification that also upregulates its activity. These two PFKFB3 protein modifications allow for flexible regulation of ATP production by glycolysis, even under simultaneous and different stresses. In fact, the constitutively active S461E PFKFB3 mutant, designed to mimic phosphorylation in response to OXPHOS inhibition, enhanced HSC reconstitution capacity after transplantation, suggesting that even if PFKFB3 is activated by one stress (in this case, proliferative), it has the activation capacity to respond to a different stress (i.e., mitochondrial). Therefore, the functions of phosphorylated and methylated forms of PFKFB3 are to some extent interchangeable, and either modification can be used to handle diverse stresses. In summary, we found that HSCs exhibit a highly dynamic range of glycolytic flux. Our study highlights glycolysis as a pivotal source of energy production in stressed HSCs, and indicates that OXPHOS, although an important source of ATP, can be uncoupled from glycolysis in steady-state HSCs without compromising ATP levels. Because multiple PFKFB3 modifications safeguard HSCs against different stresses by accelerating glycolysis, interventions targeting these might effectively induce or manage stress hematopoiesis. This study provides a platform for comprehensive and quantitative real-time analysis of ATP concentration and its dynamics in HSPCs. Our approach allows for analysis of metabolic programs in rare cells and detection of various metabolic activities within a diverse cell population, making it applicable to the analysis of various tissue systems in normal and diseased states.

## Limitations of the study

In this study, 5-FU-treated HSCs were analyzed as cell-cycling HSCs, but if more sensitive and time-saving glucose tracer analysis methods (especially after *in vivo* labeling with isotopic glucose) are developed, it may be possible to prospectively differentiate and quantitatively analyze HSC metabolism based on the cell surface antigens and cell cycle status. Although our assay uses media that mimic the BM environment, in the near future, *in vivo* GO-ATeam2 analysis will allow us to measure ATP concentrations in physiologically hypoxic BM.

## Acknowledgments

We thank E. Lamar for preparation of the manuscript, T. Kitamura for providing mVenus-p27K-mice, and N. Toyama-Sorimachi and H. Shindou for their critical reading of the manuscript. This work was supported in part by KAKENHI grants from MEXT/JSPS (JP19K17847 to H.K.; JP19K17877, JP21J01690 to D.K.; JP18H02845, JP20K21621, JP21H02957 to K.T.), AMED grants (JP22zf0127007 to M.S.; JP18ck0106444, JP18ae0201014, JP20bm0704042, JP20gm1210011 to K.T.), grants from the National Center for Global Health and Medicine (29-1015, 20A1010 to H.K.; 26-001, 21A2001 to K.T.), the Takeda Science Foundation (to D.K. and K.T.), a JB Research Grant (to D.K.), the Human Biology Microbiome Quantum Research Center (WPI-Bio2Q) supported by MEXT (to M.S.), and the MEXT Joint Usage/Research Center Program at the Advanced Medical Research Center, Yokohama City University (to K.T.).

## Author Contributions

S.W., H.K., Y. Sugiura, M.Y., K.S., Y. Sorimachi, D.K., S.K., M.O., A.N., K.M., M.H., and S.T. performed the study and analyzed data; S.F., T.M., T. Yamamoto, T. Yabushita, Y.T, G.N., H.H., S.O., N.G., T.T., A.N-I., M.S., A.I., T.S., and K.T. provided scientific advice and materials; S.W., H. K., A.N-I. and K.T. wrote the manuscript; and K.T. conceived the project and supervised the research.

