## Supplementary_Information for "Context-Dependent Modification of PFKFB3 in Hematopoietic Stem Cells Promotes Anaerobic Glycolysis and Ensures Stress Hematopoiesis"

#### Methods

##### Mice and genotyping

C57BL/6 mice (7–16 weeks old, Ly5.2$^{+}$) were purchased from Japan SLC (Shizuoka, Japan). C57BL/6 mice (Ly5.1$^{+}$) were purchased from CLEA Japan (Shizuoka, Japan). Knock-in mice harboring GO-ATeam2[1-3] in the ROSA26 locus were generated in the Yamamoto laboratory. The GO-ATeam2 mice (8–16 weeks old) were used to analyze HSPCs. Ubc-GFP reporter mice (Ubc-GFP mice) were from the Jackson Laboratory and genotyped using PCR-based assays. GO-ATeam2 mice were genotyped by PCR of tail DNA or by transdermal GFP fluorescence. The PCR protocol was as follows: 94 °C for 5 min; 34 cycles of 94 °C for 30 s, 56 °C for 30 s, 72 °C for 30 s; 72 °C for 5 min; and 4 °C hold. Primers for GO-ATeam2 or Ubc-GFP mice are listed in Table S6. mVenus-p27K$^{-}$ mice (17–20 weeks old) were provided by Kitamura Laboratory and used for cell cycle analysis[4,5]. Mice were genotyped using PCR-based assays of tail DNA or transdermal EGFP fluorescence. All mice were maintained in the animal facility at the National Center for Global Health and Medicine Research Institute under specific pathogen-free conditions and fed ad libitum. Mice were euthanized by cervical dislocation. All animal experiments were approved by the Institutional Animal Care and Use Committee (IACUC) at the National Center for Global Health and Medicine Research Institute. Both male and female mice were used.

##### Cell preparation

For C57BL/6 mice, bone marrow (BM) cells were isolated from bilateral femurs and tibiae by flushing with PBS + 2% fetal calf serum (FCS) (Gibco) using a 21-gauge needle (Terumo Corporation, Tokyo, Japan) and a 10 mL syringe (Terumo). As an exception, for U-$^{13}C_6$-labeled glucose tracer experiments using C57BL/6 mice, BM was flushed with PBS + 0.1% bovine serum albumin (BSA, Cat# A4503). The BM plug was dispersed by refluxing through the needle, and the suspension was centrifuged 680 × *g* for 5 min at 4 °C. Cells were lysed with lysis buffer (0.17 M $NH_4Cl$, 1 mM EDTA, 10 mM $NaHCO_3$) at room temperature (RT) for 5 min, washed with two volumes PBS + 2% FCS (or PBS + 0.1% BSA for tracer experiments), and centrifuged at 680 × *g* for 5 min at 4 °C. Cells were resuspended in PBS + 2% FCS (or PBS + 0.1% BSA for tracer experiments) and filtered through 40 µm nylon mesh (BD Biosciences). Cells were again centrifuged 680 × *g* for 5 min at 4 °C and treated with anti-CD16/32 antibody for Fc-receptor block (2 µL/mouse; BD Biosciences, Cat# 553152) for 10 min at

4 °C. Anti-c-Kit magnetic beads (Miltenyi Biotec, Bergisch Gladbach, Germany, Cat# 130-091-224) were added at a 1:5 v/v ratio for 15 min at 4 °C. After removing the antibody with two PBS + 2% FCS (or PBS + 0.1% BSA for tracer experiments) washes, c-Kit-positive cells were isolated using Auto-MACS Pro (Miltenyi Biotec) with the Possel-s or Possel-d2 program. Isolated cells were centrifuged once at 340 × *g* for 5 min and stained with an antibody cocktail for flow cytometry.

For analysis of the GO-ATeam2 hematopoietic cells, BM from GO-ATeam2 mice was flushed with PBS + 0.1% BSA to minimize exposure to nutrients in FCS. Hemolysis, centrifugation, filtering, and Fc receptor blocking were performed in the same manner as for cell preparation using C57BL/6 mice. Cells were stained for 30 min with an antibody cocktail at 4 °C and then washed and suspended in 1000 µL PBS + 0.1% BSA and centrifuged at 340 × *g* at 4 °C for 5 min. Supernatants were discarded in preparation for flow cytometry.

**Flow cytometry and cell sorting**

Murine hematopoietic stem and progenitor fractions were labeled as follows: To stain cells from C57BL/6 mice, lineage (Lin) markers (CD4, CD8a, Gr-1, Mac-1, Ter-119, B220)-PerCP-Cy5.5 (BD Biosciences for CD4 (Cat# 550954), Gr-1 (Cat# 552093), Mac-1 (Cat# 550993), B220 (Cat# 552771) and BioLegend for CD8a (Cat# 100734) and Ter-119 (Cat# 116228) antibodies), c-Kit-APC-Cy7 (BioLegend, Cat# 105826), Sca-1-PE-Cy7 (BioLegend, Cat# 122514), CD150-PE (BioLegend, Cat# 115904), CD48-FITC (BioLegend, Cat# 103404), and Flt3-APC (BioLegend, Cat# 135310) were used. For HSC collection five days after 5-FU administration (intraperitoneally or intravenously), Mac-1 antibody was excluded from the antibody cocktail1, and the LSK gate was expanded to include c-Kit-high to -dim Lin$^-$ cells to include functional HSCs early after 5-FU administration as previously reported[6 7]. We did not expand the LSK gate at any time other than five days after 5-FU administration. When sorting or analyzing EPCR$^+$CD150$^+$CD48$^-$LSK cells from C57BL/6 mice or mVenus-p27K- mice, CD150-BV421 (BioLegend, Cat# 115926), CD48-APC (BioLegend, Cat# 103412), and EPCR-PE (Biolegend, Cat# 141503) were used in addition to LSK for staining, and FLT3 staining was excluded. To stain cells from GO-ATeam2 mice or C57BL/6 mice for the 2-NBDG assay or homing assay using Ubc-GFP mice, lineage markers (CD4, CD8a, Gr-1, Mac-1, Ter-119, B220)-PerCP-Cy5.5, c-Kit-APC-Cy7, Sca-1-PE-Cy7, CD150-BV421, and CD48-APC were used. In the analysis using GO-ATeam2 mice, Flt3 was not used to define HSCs because the fluorescence of the FRET sensor (EGFP, mKO) limits the available fluorochromes for surface marker staining. In analysis using the AMPK inhibitor dorsomorphin (Cayman Chemical, Cat# 21207), CD150-APC (BioLegend, Cat# 115910) and CD48-AlexaFluor700 (BioLegend, Cat# 103426) were used to stain LSK-SLAM to eliminate effects of dorsomorphin fluorescence on cell staining. Cells were resuspended in 0.5–2 mL of PBS + 2% FCS + 0.1% propidium iodide (PI) (Invitrogen, Cat# P3566) (for C57BL/6 mice) or PBS + 0.1% BSA (for GO-ATeam2 mice) and sorted using the FACSAria IIIu Cell Sorter (BD Biosciences) into RPMI1640 (without glucose) (Nacalai Tesque, Cat# 09892-15) containing 4% w/v BSA or GO-ATeam2 basal medium (Ba-M, Table S1 with 4% w/v BSA) (custom made by Gmep Inc.). Murine HSCs were defined as $CD150^{+}CD48^{-}Flt3^{-}LSK$ (for C57BL/6 mice) or $CD150^{+}CD48^{-}LSK$ (for GO-ATeam2 mice and mVenus-p27K- mice, and when EPCR was included in the antibody cocktail against C57BL/6 mice) cells. MPPs were defined as $CD150^{-}CD48^{+}Flt3^{-}$ LSK (for C57BL/6 mice) or $CD150^{-}CD48^{+}LSK$ (for GO-ATeam2 mice) cells. Among myeloid progenitors (MyPs), GMPs/MEPs/CMPs were defined as follows: GMPs ($CD16/32^{+}$ $CD34^{+}$), MEPs ($CD16/32^{-}$ $CD34^{+}$), and CMPs ($CD16/32^{-}$ $CD34^{+}$). CLPs were defined as $Lin^{-}Sca\text{-}1^{low}c\text{-}Kit^{low}Flt3^{+}IL7R\alpha^{+}$ cells. Data were analyzed using FlowJo™ V10 (Tree Star) software.

#### Intracellular staining for phosphorylated Rb (pRb)

$EPCR^{+}$ or $EPCR^{-}$ LSK-SLAM cells from PBS- or 5-FU-treated C57BL/6 mice were purified separately (see "**Flow cytometry and cell sorting**" for details). Anti-phospho-Rb (Ser807/811) antibody (CST, Cat# 8516T) was used as the primary antibody and Anti-rabbit IgG (H+L), F(ab') Fragment (Alexa Fluor488 Conjugate) (CST, Cat# 4412) was used as the secondary antibody. Fixation and permeabilization were performed according to the manufacturer protocol. pRb and DNA content (stained with PI) were analyzed by flow cytometry.

#### Analysis of mVenus-p27K-mouse-derived BM cells

Surface-marker-stained BM mononuclear cells (MNCs) (see "**Flow cytometry and cell sorting**" for details) were analyzed by flow cytometry to determine the frequency of G0 marker positivity for $EPCR^{+}$ or $EPCR^{-}$ $CD150^{+}CD48^{-}LSK$ or progenitor cells.

#### Seahorse flux analyzer

The extracellular acidification rate (ECAR) and oxygen consumption rate (OCR) were measured using a Seahorse XFe96 extracellular flux analyzer according to the manufacturer's instructions (Agilent Technologies). Briefly, sorted cells were dispensed to culture plates pre-coated with Cell-Tak (Corning) and then the media was replaced with pre-warmed XF-DMEM medium (Agilent) supplemented with 10 mM glucose, 1 mM pyruvate, and 2 mM glutamine, followed by centrifugation at 200 × *g* for 5 min. OCR and ECAR were measured at baseline and again after sequential addition of respiratory inhibitors at final concentrations of 1 μM oligomycin (an inhibitor of ATP synthase), 2 μM FCCP (an uncoupling agent of mitochondrial respiration), 0.5 μM rotenone/antimycin (an inhibitor of mitochondrial complex I/III) and 50 mM 2-deoxy-D-glucose (an inhibitor of glycolysis). The experiment was performed by dispensing 75,000 HSCs (PBS or 5-FU treated) or MyPs per well.

**2-NBDG assay**

For the *in vitro* 2-NBDG assay, sorted HSCs were exposed to 200 μM 2-NBDG (Cayman Chemical, Cat# 11046) for 30 min. HSCs were then centrifuged at 340 × *g* at 4 °C for 5 min, and the supernatant was removed. The uptake of 2-NBDG was measured using FACS Aria IIIu. As a negative control, HSCs were simultaneously exposed to 54 μg/ml phloretin or 20 μg/ml cytochalasin B with 2-NBDG. For the *in vivo* 2-NBDG assay, C57BL/6 mice treated with PBS or 5-FU were subjected to an *in vivo* 2-NBDG assay as reported by Jun et al.[8]. Mice received a bolus dose of 375 μg 2-NBDG intravenously and were euthanized by cervical dislocation after 1 h. Mice were immediately placed on ice, and all subsequent cell preparation processes were performed while the cells were chilled on ice. The 2-NBDG positive cell fraction was detected by flow cytometry.

**Conversion of GO-ATeam2 fluorescence to ATP concentration**

The GO-ATeam2 knock-in mice were reported by Yamamoto et al [3]. Briefly, we used a CAG promoter-based knock-in strategy targeting the *Rosa26* locus to generate GO-ATeam2 knock-in mice. A study presenting the significance of measuring the absolute concentration of ATP at the single-cell level is currently in preparation for submission, but briefly, the FRET efficiency was converted to the absolute concentration of ATP using the following method (Watanuki et al., *in preparation*). To permeabilize BM cells, α-hemolysin stock solution (Sigma-Aldrich, St. Louis, MO, USA) was diluted in permeabilization buffer (140 mM KCl (Wako, Cat# 163-03545), 6 mM NaCl (Wako, Cat# 191-01665), 0.1 mM EGTA (Wako, Cat# QB-6401), and 10 mM HEPES (Wako, Cat# 342-01375) [pH 7.4]) to a final concentration of 50 μg/mL α-hemolysin. GO-ATeam2-knock-in BMMNCs were added to the buffer and permeabilized for 30 min at 37 °C under 5% $CO_2$. To calibrate ATP concentration, calibration buffer (140 mM KCl, 6 mM NaCl, 0.5 mM MgCl2 (Wako, Cat# 136-03995), and 10 mM HEPES [pH 7.4]) and Mg-ATP stock solution (Sigma-Aldrich, Cat# A9187) were prepared. After washing GO-ATeam2-knock-in BMMNCs with calibration buffer, fresh calibration buffer without ATP was added. Mg-ATP was gradually added to increase ATP concentration in the cell suspension, and

FRET values of the GO-ATeam2 biosensor at defined ATP concentrations were analyzed by flow cytometry. The FRET value (relative ratio of FRET to EGFP fluorescence intensities) was calculated by the following equation.

$$FRET\ value = \frac{\text{Fluorescence of FRET}}{\text{Fluorescence of EGFP}} \ldots (1)$$

The excitation wavelength of FRET and EGFP was set at 488 nm.

The FRET value was then fitted to Hill's formula[9] as a function of ATP concentration:

$$\theta = \frac{[L]^n}{[K_A]^n + [L]^n} \ldots (2)$$

where $\theta$ is the original percentage of receptor proteins occupied by the ligand, $[L]$ is the free (unbound) ligand concentration, $K_A$ is the concentration of ligand at half saturation, and $n$ is Hill's coefficient.

Equation (2) was transformed as $\log\left(\frac{\theta}{1-\theta}\right) = n \log[L] - n \log K_A \ldots (3)$

such that $\theta$ could be expressed by the FRET value as follows:

$$\theta = \frac{FRET\ value - 1.4}{6} \ldots (4)$$

We estimated parameters $n$ and $K_A$ by fitting observed FRET values to the linear regression model represented in equation (3). In our experiment, $n = 3.1234$, and $K_A = 0.84699$. Using these parameters, cellular ATP concentration, $[L]$, was estimated.

**Time-course analysis of FRET values**

GO-ATeam2 is a ratiometric biosensor that monitors ATP concentration through Förster resonance energy transfer (FRET) from EGFP to the monomeric version of Kusabira Orange (mKO), regardless of the sensor expression levels[2]. Surface-marker-stained BMMNCs from GO-ATeam2 mice were dispensed into a basal medium (Ba-M) containing minimal salts, vitamins, and buffers (HEPES and sodium bicarbonate), but no glucose or mitochondrial substrates (Table S1), or into a medium containing mitochondrial substrates, pyruvate, lactate, fatty acids, and amino acids, but no glucose (PLFA medium). Depending on the experiment, fresh and surface marker-stained BMMNCs obtained from mice two, five, or 14 days after intraperitoneal administration of PBS or 5-FU were dispensed into a Ba-M or PLFA medium. The FRET/EGFP ratio data was imported continuously during analysis in a real-time manner using the BD FACSAria IIIu under ambient pressure. Depending on their purpose, experiments were conducted in the presence or absence of various nutrients or metabolic modulators (Supplemental Figure 4A). For this platform, 2 mL of Ba-M or PLFA medium per tube was pre-

saturated with 1% $O_2$/5% $CO_2$/94% $N_2$ to stabilize ATP levels of BMMNCs (Supplemental Figure 4B) and mimic the hypoxic BM environment; when medium was not pre-saturated, ATP concentrations rapidly decreased, even in the presence of glucose, pyruvate, or lactate (Supplemental Figure 4C).
To reduce the effect of autofluorescence as much as possible, the top 40–50% of EGFP and FRET fractions of MFI were used in the analysis (MFI > 1000 for EGFP and FRET). Then, data reporting EGFP and FRET fluorescence values in individual cells from each gating (e.g. HSCs, MPPs) were extracted along with time course data. Relevant nutrients and inhibitors were added to medium with samples for analysis. Data acquired by the FACSAria IIIu device and retrieved as FCS files were analyzed by the flowCore package in R software. The FRET/EGFP ratio of each set of single cells was fitted to a generalized additive model using the 'gam' function in the 'mgcv' package with 's', a spline-based smoothing function, in default settings as a function of time, then smoothened using the 'predict' function. Pseudocolor plots of the FRET/EGFP ratio were created using the 'kde2d function'. If needed, fitted data were converted to ATP concentration using the model described above.

To compare changes in ATP concentrations in PBS- and 5-FU-treated groups, we corrected differences in baseline ATP concentrations by multiplying all data from the PBS-treated group by the following value: ATP concentration at 0 s in the 5-FU group/ATP concentration at 0 s in the PBS group.

**Ki67/Hoechst staining**

Ki67 (BD Biosciences, Cat# 558617) and Hoechst 33432 (Invitrogen, Cat# H3570) were used for cell cycle analysis of fixed cells from C57BL/6 mice. A total of $4\times10^6$ BMMNCs/sample were stained with anti-CD150-APC, anti-CD48-FITC, anti-lineage (CD4, CD8a, Gr-1, Mac-1, Ter-119, B220)-PerCP-Cy5.5, anti-c-Kit-APC-Cy7, and anti-Sca-1-PE-Cy7 antibodies. To stain samples after 5-FU treatment, Mac-1 was excluded from the antibody cocktail. Stained samples were centrifuged at 340 × *g* and 4 °C for 5 min. To analyze HSCs derived from mice after *in vivo* 2-NBDG administration, anti-CD150-PE and anti-CD48-Alexa700 antibodies were alternatively used to sort and purify HSCs with high or low NBDG uptake. These HSCs were then subjected to Ki67/Hoechst staining. Next, 250 µL of BD Cytofix/Cytoperm (BD Biosciences, Cat# 555028) was added, and samples were incubated for 20 min at 4 °C for fixation. Fixed cells were centrifuged and washed twice at 340 × *g* at 4 °C, with 1 mL BD Perm/Wash buffer (BD Biosciences, Cat# 554723) diluted 10-fold. Each sample was stained with 10 µL of Ki67-Alexa555 or Ki67-eFlour660 (for 2-NBDG stained HSC) antibodies for 1 h at RT, shaded from light. Ki67-stained cells were centrifuged and washed twice at 340 × *g* and 4 °C with PBS. Samples were resuspended in 500 µL of PBS + 10 µg/mL Hoechst 33432, filtered, and analyzed with the BD FACSAria IIIu instrument.

**FAOBlue assay**

Surface marker-stained BMMNCs from PBS- or 5-FU-treated mice were dispensed at $3 \times 10^5$ cells in 500 µL Ba-M, which had been pre-saturated for 48 h under 1% $O_2$ and 5% $CO_2$ conditions and contained 200 mg/dL glucose and 50 µM verapamil. These cells were then exposed to 5 µM FAOBlue (Funakoshi) for 15 min. As a negative control, BMMNCs were exposed to 100 µM etomoxir simultaneously with FAOBlue. The FAOBlue-stained BMMNCs were then centrifuged at 340 × *g* for 5 min at 4°C, and the supernatant was discarded. The fluorescence of FAOBlue was excited at a wavelength of 405 nm and detected in the V 450/50 channel. After analysis, the HSC fraction data were extracted.

**Analysis of peripheral blood and BM chimerism**

Periorbitally collected peripheral blood from BMT recipients was centrifuged for 3 min at 340 × *g* and the supernatant discarded. Samples were subjected to hemolysis with 1000 µL of 0.17 M $NH_4Cl$ for 40–50 min and centrifuged at 340 × *g* for 5 min. The supernatant was discarded, and samples were again subjected to hemolysis with 1000 µL of 0.17 M $NH_4Cl$ for 10–20 min. Samples were centrifuged again at 340 × *g* for 5 min and the supernatant was discarded. Pellets were then resuspended in 50 µL PBS and 0.3 µL Fc receptor block and incubated at 4 °C for 5 min. Surface antigen staining was performed using the following antibody panel: Gr-1-PE-Cy7 (BioLegend, Cat# 108416), Mac-1-PE-Cy7 (BioLegend, Cat# 101216), B220-APC (BioLegend, Cat# 103212), CD4-PerCP-Cy5.5, CD8a-PerCP-Cy5.5, CD45.1-PE (BD Biosciences, Cat# 553776), and CD45.2-FITC (BD Biosciences, Cat# 553772). An antibody cocktail was prepared by mixing 0.3 µL of each antibody. The frequency (%) of donor-derived cells was calculated as follows:

The frequency (%) of donor-derived cells = 100 × Donor-derived ($Ly5.2^+Ly5.1^-$) cells (%) / (Donor-derived cells [%] + Competitor- or recipient-derived [$Ly5.2^-Ly5.1^+$] cells [%]].

Myeloid, B, and T cells were identified by $Gr\text{-}1^+$ or $Mac\text{-}1^+$, $B220^+$, or $CD4^+$ or $CD8^+$, respectively.

Four months after BM transplant, the frequency of donor-derived cells in BM was determined using one femur and tibia per recipient. Anti-CD150-BV421, anti-CD48-PE (BD Biosciences, Cat# 557485), anti-lineage (CD4, CD8a, Gr-1, Mac-1, Ter-119, B220)-PerCP-Cy5.5, anti-c-Kit-APC-Cy7, anti-Sca-1-PE-Cy7, anti-Ly5.1-Alexa-Fluor700 (BioLegend, Cat# 110724), and anti-Ly5.2-FITC antibodies were used for surface antigen detection. An antibody cocktail was prepared by mixing 1 µL of each antibody.

#### Comparison of metabolite levels before and after sorting

c-Kit-positive cells were isolated using Auto-MACS Pro (Miltenyi Biotec) with the Possel-s or Possel-d2 program as described above (see "**Cell preparation**" for details). Isolated cells were counted, and $1 \times 10^5$ viable cells were dispensed into methanol containing an internal standard as a pre-sorting cell sample and stored at −80 °C until IC-MS analysis. To the isolated cell suspension, 0.1% PI was added and samples were sorted using the FACS Aria IIIu. A total of $1 \times 10^5$ viable cells ($PI^-$ cells) were sorted directly into methanol containing an internal standard as a post-sorting cell sample and stored at −80 °C until IC-MS analysis. The detected metabolites were quantified based on calibration curve data (see "**Ion chromatography mass spectrometry (IC-MS) analysis**" in "**Methods**" for details).

#### Preparation and storage of *in vitro* U-$^{13}C_6$-glucose tracer samples

For tracer analysis, C57BL/6 mice were euthanized to obtain 25,000–50,000 cells per sample of each fraction (HSC, MPP, GMP, CLP) from BM using the FACSAria IIIu instrument. Numbers of mice used to obtain each fraction were as follows: 30–35 each for steady state HSCs and MPPs, 60–65 each for 5-FU treated HSCs, 10 each for GMPs and CLPs. In addition, bone and BM cells were chilled by placing dishes and tubes on ice during the cell preparation process; samples were washed with ice-cold buffer throughout the entire process before cell sorting. Experiments and experimental manipulations regarding the sampling of mouse femurs and tibias were also performed in the shortest amount of time possible by skilled personnel. Cells were sorted in 0.1% BSA + PBS and sorted cells were centrifuged at 340 × *g* and 4 °C for 5 min. After discarding the supernatant, cells were added to 1 mL pre-saturated (under 1% $O_2$ and 5% $CO_2$) GO-ATeam2 Ba-M + 0.1% BSA + 200 mg/dL U-$^{13}C_6$- (Sigma-Aldrich, Cat# 389374) or U-$^{12}C_6$-glucose and incubated 10 or 30 min. If the process of pre-saturation was omitted, ATP levels dropped rapidly within a short time (Supplementary Figure 5G).When using oligomycin (1 μM) (Cell Signaling Technology, Cat# 9996), exposure time was set to 10 min. Samples were then immediately centrifuged at 1000 × *g* and 4 °C for 3 min. After discarding supernatants, cells were frozen and stored at −80 °C.

#### Preparation and storage of *in vivo* U-$^{13}C_6$-glucose tracer samples

U-$^{13}C_6$-glucose administration to C57BL/6 mice was performed based on the methods of Jun et al.[8], with some modifications. Mice were intraperitoneally administered medetomidine hydrochloride, midazolam, and butorphanol tartrate at 0.75 mg/kg, 4 mg/kg, and 5 mg/kg, respectively. After anesthesia, mice were kept warm on a hot plate set at 37 °C while a 27-gauge needle was placed in the external tail vein and U-$^{13}C_6$-glucose was continuously administered. The dose and duration of U-$^{13}C_6$-

glucose administration followed Jun et al.[8], and 0.4125 mg/g body mass was administered in 1 min, followed by 0.008 mg/g body mass per minute for 3 h. After U-$^{13}C_6$-glucose administration, mice were euthanized by cervical dislocation and immediately placed on ice. For *in vivo* tracer analysis, BMMNCs from the bilateral femur, tibia, pelvis, and sternum of each mouse were used to prepare sufficient numbers of HSCs, and pre-chilled 0.1% BSA+PBS was used for BM flushing and washing. HSCs were directly sorted in methanol and stored at −80 °C until IC-MS analysis. A total of 1–3 × $10^4$ HSCs were purified from one or two mice in the PBS group and from two or three mice in the 5-FU group.

When generating the heat map of labeling rates in each metabolite, 1 was added as a pseudo number to the labeling rate of all metabolites. When calculating the total amount of $^{13}C$ labeled metabolites for each pathway, metabolites other than M+0 were summed in each metabolite.

**Metabolite extraction**

Frozen samples were mixed with 500 µL methanol containing internal standards and sonicated for 10 s. Then, 200 µL dd$H_2O$ (Invitrogen, Cat# 10977-015) and 400 µL chloroform (Nacalai tesque, Cat# 08402-55) were added and samples were centrifuged at 10000 × *g* and 4 °C for 3 min. The aqueous phase was transferred to an Amicon ultrafiltration system (Human Metabolome Technologies, Inc., Cat# UFC3LCCNB-HMT) and centrifuged at 9100 × *g* and 4 °C for 3 h. Filtered samples were analyzed by IC-MS.

**Ion chromatography mass spectrometry (IC-MS) analysis**

For metabolome analysis focused on glycolytic metabolites and nucleotides, anionic metabolites were measured using an orbitrap-type MS (Q-Exactive Focus; Thermo Fisher Scientific, Waltham, MA, USA) connected to a high-performance IC system (ICS-5000+, Thermo Fisher Scientific), enabling highly selective and sensitive metabolite quantification owing to the IC-separation and Fourier Transfer MS principle[10]. The IC instrument was equipped with an anion electrolytic suppressor (Dionex AERS 500; Thermo Fisher Scientific) to convert the potassium hydroxide gradient into pure water before the sample entered the mass spectrometer. Separation was performed using a Dionex IonPac AS11-HC-4 µm IC column (Thermo Fisher Scientific). The IC flow rate was 0.25 mL/min supplemented post-column with a 0.18 mL/min makeup flow of MeOH. The potassium hydroxide gradient conditions for IC separation were as follows: 1–100 mM (0–40 min), 100 mM (40–50 min), and 1 mM (50.1–60 min), with a column temperature of 30 °C. The Q-Exactive Focus mass spectrometer was operated under the ESI negative mode for all detections. A full mass scan (*m/z*

70−900) was performed at a resolution of 70,000. The automatic gain control target was set at $3 \times 10^6$ ions, and the maximum ion injection time was 100 ms. Source ionization parameters were optimized with a spray voltage of 3 kV, and other parameters were as follows: transfer temperature, 320 °C; S-lens level, 50; heater temperature, 300 °C; sheath gas, 36; and aux gas, 10. Metabolite amounts were quantified from calibration curve data generated based on peak areas and respective metabolite amounts.

**Quantitative $^{13}$C-MFA with OpenMebius**

OpenMebius (Open source software for $^{13}$C-MFA) provides the platform to simulate isotope labeling enrichment from a user-defined metabolic model setup worksheet developed in MATLAB (MathWorks, Natick, MA, USA)[11]. Quantitative $^{13}$C-MFA was performed according to a manual prepared by the software developer (http://www-shimizu.ist.osaka-u.ac.jp/hp/en/software/OpenMebius.html), but some metabolic model modifications were made to more faithfully reflect our measured data. Specifically, the model was modified to include (a) the conversion of pyruvate to lactate catalyzed by lactate dehydrogenase, (b) the formation of citrate from acetyl CoA and oxaloacetate catalyzed by citrate synthase, (c) the synthesis of alpha-ketoglutarate from citrate catalyzed by aconitase and isocitrate dehydrogenase, and (d) the synthesis of fumarate from succinate by succinate dehydrogenase. Reactions with pyruvate formate lyase performed by *Escherichia coli*, *Streptococcus spp.*, and ethanol fermentation of acetyl CoA were excluded from the default metabolic network sheet.

The lactate efflux values in $^{13}$C-MFA were determined using the following trial and error method. First, various values (0–100) were entered as candidate lactate efflux values and simulations were run to determine the optimal lactate efflux. When the lactate efflux value was set low (below 50), either the simulation could not be run and an error occurred, or the simulation resulted in the glycolytic system progressing in the opposite direction. These results suggested that the appropriate solution was not obtained because the lactate efflux was unnatural compared to the level of glycolytic metabolites. This was validated by experimental data showing that isotopic labeling rates for most glycolytic metabolites were close to 100% at short labeling times (Supplementary Figure 2C). Therefore, we ran the simulation with a higher lactate efflux value. Finally, we set the lactate efflux to 65, which yielded reasonably satisfactory results for nearly 100% labeling of glycolytic and PPP metabolites in PBS- or DMSO-treated HSCs.

The rate of lactate efflux 5-FU-treated HSCs with the rate of glucose uptake set to 100 was defined using the following equation, with the flux in stationary phase HSC set to 65:

65×(Percentage of glycolytic metabolites labeled with $^{13}C$ in the total $^{13}C$-labelled metabolites [5-FU-treated HSCs])/(Percentage of glycolytic metabolites labeled with $^{13}C$ in the total $^{13}C$-labelled metabolites [PBS-treated HSC])

In the metabolic flux measurements of HSCs under mitochondrial stress, the lactate efflux determined by the above method exceeded the maximum value that could be modeled (85>), so we decreased the lactate efflux flux by 5 and adopted the maximum value, 80, at which modeling became possible. For values of efflux other than those of lactate efflux flux, the values specified by the OpenMebius manual were used to eliminate arbitrary factors as much as possible.

The metabolic substrate used for labeling was set to 100% U-$^{13}C_6$ glucose. Metabolites used in the analysis included the first intermediate metabolite produced when U-$^{13}C_6$ glucose is metabolized (e.g., G6P or F6P with all carbons labeled, the labeled metabolite of the first cycle of the TCA cycle) and the unlabeled metabolite that was measured. Some of the labeled metabolites in the TCA cycle (e.g., citrate [M2]) and erythrose 4-phosphate (M4) in PPP were detected with non-negligible amounts of natural isotopes (>5% even when labeled with U-$^{12}C_6$ glucose compared to U-$^{13}C_6$ glucose). The presence of such natural isotopes may result in overestimation of the amount of increased labeling with U-$^{13}C_6$ glucose. In such cases, the amount of natural isotope detected when labeled with U-$^{12}C_6$ glucose was subtracted from the amount of labeled metabolite detected with U-$^{13}C_6$ glucose. If the resulting true labeled isotope abundance was negative, the labeled amount was modeled as zero. When analyzing in MATLAB, the number of modeling cycles was set to 100, and the iteration time was set to a maximum of 2000 cycles.

**Luminometric ATP measurement**

HSCs were sorted from C57BL/6 mice treated with PBS or 5-FU and dispensed into pre-saturated GO-ATeam2 medium with 0.1% BSA in a 1%$O_2$/5%$CO_2$ incubator. HSCs were then exposed to 15 μM of PFKFB3 inhibitor (AZ PFKFB3 26) or DMSO and placed in a 1%$O_2$/5%$CO_2$ incubator for 10 min. Cells were centrifuged at 4 °C and 340 × *g* and the supernatant was removed. ATP measurements were performed according to manufacturer instructions using Cell ATP Assay Reagent Ver. 2 (Toyo B- Net Corporation). The amount of ATP per cell was calculated by dividing the amount of ATP detected by the number of cells used for analysis.

**Apoptosis assay of HSC after 2-DG or oligomycin treatment**

Purified C57B6/J mouse-derived HSCs were exposed to 2-DG (50 mM) and oligomycin (1 μM) in pre-saturated 0.1% BSA+GO-ATeam2 medium under 1% $O_2$/5% $CO_2$ conditions for 10 min and subjected to apoptosis assay using the PE Annexin V Apoptosis Detection Kit I (BD Biosciences, Cat# 559763) according to manufacturer instructions.

**CRISPR/Cas9 knockout (KO) of *Pfkfb3***

Target sequences of single guide RNA (sgRNA) were provided in a previous report[12] and identified using the web tool GenScript (https://www.genscript.com) for *Pfkfb3*. sgRNAs were synthesized using a CUGA7 gRNA Synthesis Kit (Nippon Gene, Tokyo, Japan, Cat# 314-08691) following manufacturer instructions, diluted to 1.5 µg/µL, and cryopreserved at −80 °C until use. $CD150^+CD48^-Flt3^-$ LSK cells sorted by FACSAria IIIu were cultured in SF-O3 medium supplemented with stem cell factor (SCF) (50 ng/mL) (Peprotech, Cat# 250-03) and thrombopoietin (TPO) (Peprotech, Cat# 300-18) (50 ng/mL) (S50T50 medium) and incubated under 20% $O_2$/5% $CO_2$ conditions for 16–24 h, enabling subsequent HSC-specific gene editing with the CRISPR-Cas9 system. Ribonucleoprotein complex preparation and electroporation were conducted as previously reported [13]. Briefly, 3 µg Cas9 protein (TrueCut Cas9 Protein v2, Thermo Fisher Scientific, Cat# A36496) plus 3 µg of sgRNA were incubated in Buffet T (Invitrogen, Cat# MPK10096) for 20 min at RT in a volume 6 µL. Cultured cells were resuspended in 30 µL Buffer T and added to ribonucleoprotein at a total volume of 36 µL. Cells were electroporated using the Neon Transfection System (Thermo Fisher Scientific) at 1700 V for 20 ms with one pulse. The cell suspension was transferred to S50T50 medium and cultured under 20% $O_2$/5% $CO_2$ conditions. To evaluate gene editing efficiency, genomic DNA from LSK cells was extracted using the NucleoSpin system (Macherey-Nagel, Dürin, Germany) 2–3 d after electroporation. PCR was performed using the following settings: 95 °C for 2 min; 35 cycles of 95 °C for 30 s, 60 °C for 30 s, and 72 °C for 30 s; followed by final extension at 72 °C for 5 min. PCR products were purified using Wizard SV Gel and the PCR Clean-Up System (Promega Corporation, Madison, WI, USA, Cat# A9281) following manufacturer instructions. A tracking of indels by decomposition (TIDE) assay[14] or inference of CRISPR edits analysis[15] was performed to analyze the sequence data of each PCR product obtained by Sanger sequencing. Among five sgRNAs, Pfkfb3-sg1 displayed the best editing efficiency and was used for subsequent transplant and culture experiments.

**BM transplant of *Pfkfb3*-KO HSCs**

Either *Rosa26* (control) or *Pfkfb3* sequences in HSCs were targeted using CRISPR/Cas9. After electroporation, HSCs were incubated for 2–3 h in S50T50 medium under 5%$CO_2$/20%$O_2$ conditions, and then counted using a TC10 Automated Cell Counter (Bio-Rad Laboratories, Inc., Hercules, CA,

USA). Subsequently, 500 gene-edited HSCs together with 2 × $10^6$ BM cells from Ly5.1 congenic mice were transplanted retro-orbitally into lethally (9.5Gy using MBR-1520R with a 125 kV 10 mA, 0.5 mm Al, 0.2 mm Cu filter)-irradiated Ly5.1 mice. During *Pfkfb3* KO using the vector-free CRISPR-Cas9 system, the KO efficiency was not 100%, so the transplanted cells were a mixture of *Pfkfb3*-KO cells and wild-type cells. Therefore, after 2, 8, and 16 weeks, peripheral blood was collected and donor-derived chimerism was assessed by a TIDE assay based on a recent study by Shiroshita et al.[16]. The following oligonucleotides for sgRNA synthesis and primers for post-knockout genomic PCR were used.

For *Rosa* region KO:

sgRNA target: 5′-ACTCCAGTCTTTCTAGAAGA-3′
Forward primer 1: 5′-CCAAAGTCGCTCTGAGTTGTTATCAGT-3′
Reverse primer 1: 5′-GGAGCGGGAGAAATGGATATGAAG-3′
Forward primer 2: 5′-CCAAAGTCGCTCTGAGTTGTTATCAGT-3′
Reverse primer 2: 5′-GGAGCGGGAGAAATGGATATGAAG-3′
Sequence primer: 5′-ACATAGTCTAACTCGCGACAC-3′

For *Pfkfb3* KO:

sgRNA target: 5′-GTTGGTCAGCTTCGGCCCAC-3
Forward primer: 5′-AATTGTGTAGCACAGGATCACC-3′
Reverse primer: 5′-GCCACTAAAGGAAGGCTAGTTAC-3′
Sequence primer: 5′-CTCAATCTTCCCGAGTCTGTCTC-3′

For *CD45* KO:

sgRNA target: 5′-GGGTTTGTGGCTCAAACTTC-3′
Forward primer: 5′-AGAAGCCATTGCACTGACTTTG-3′
Reverse primer: 5′-GTGTGATCTTTCCCCGAAACAT-3′
Sequence primer: 5′-CTGCAAAGAGGACCCTTTACAGT-3′

To calculate the KO efficiency of the *Rosa* locus, primer 1 or primer 2 was used for PCR amplification.

**_Pfkfb3_ overexpression in GO-ATeam2$^+$ HSCs and time-course analysis of FRET values**

cDNA encoding *Pfkfb3* was subcloned into pMY-IRES-hCD8 upstream of IRES-hCD8. To produce a recombinant retrovirus, plasmid DNA was transfected into Plat-E cells using FuGENE® HD Transfection Reagent (Promega, Cat# E2311). Cell supernatants were then used to transduce GO-ATeam2$^+$ HSCs pre-cultured with SCF and TPO for 16 h. At 48 h post-transduction, surface-marker-stained, retrovirally *pfkfb3*-overexpressed GO-ATeam2$^+$ cells were used for time-course analysis of

FRET values as described above subsection "**Time-course analysis of FRET values**." Cells transduced with pMY-IRES-hCD8 retrovirus served as controls. Transduced cells were stained with the following antibody panel: lineage markers (CD4, CD8a, Gr-1, Mac-1, Ter-119, B220)-PerCP-Cy5.5, c-Kit-APC-Cy7, Sca-1-PE-Cy7, CD150-BV421, CD48-BV510 (BD Biosciences, Cat# 563536), and hCD8-APC (BioLegend, Cat# 980904). FRET value data for hCD8-positive cells were used for subsequent conversion to ATP concentration.

**_Pfkfb3_/_Pfkfb3_CA overexpression in HSCs and BMT**

cDNA encoding *Pfkfb3* or the constitutively active S461E *Pfkfb3* mutant (*Pfkfb3*CA)[17] was subcloned into pMY-IRES-hCD8 upstream of IRES-hCD8 or into pMY-IRES-EGFP upstream of IRES-EGFP[18], respectively. To produce a recombinant retrovirus, plasmid DNA was transfected into Plat-E cells using the FuGENE® HD Transfection Reagent. Cell supernatants containing virus were then filtered with Millex-HV Syringe Filter Unit (0.45 µm, PVDF, 33 mm, gamma sterilized, Millipore) and used to transduce Ly5.1+ HSCs pre-cultured in SCF and TPO for 16 h.

At 48 h post-transduction, 2000 transduced $GFP^+$ cells were sorted and transplanted, together with $4 \times 10^5$ BMMNCs from C57BL/6-Ly5.2 mice, into lethally (9.5Gy using MBR-1520R with a 125 kV 10 mA, 0.5 mm Al, 0.2 mm Cu filter)-irradiated C57BL/6-Ly5.2 mice. Cells transduced with pMY-IRES-EGFP retrovirus served as controls. After 1–4 months, peripheral blood was collected and donor-derived chimerism was analyzed by flow cytometry. The frequency (%) of donor-derived cells was calculated as follows:

100 × Donor-derived ($Ly5.2^-Ly5.1^+$) cells (%) / (Donor-derived cells [%] + Competitor- or recipient-derived [$Ly5.2^+Ly5.1^-$] cells [%])

**Knockout and overexpression of *Pfkfb3* in HSPC and non-competitive BMT**

PFKFB3 was knocked out and overexpressed in FACS-sorted $Lin^-Sca\text{-}1^+c\text{-}Kit^+$ and $Ly5.2^+$ cells, respectively. Methods were partially modified from those described in the "**CRISPR/Cas9 KO of *Pfkfb3***" and "**_Pfkfb3_/_Pfkfb3_CA overexpression in HSCs and BMT**" sections.

For KO of *Pfkfb3*, triple-gRNA purchased from Synthego (Redwood City, CA, USA) was used. After gene editing, $Ly5.2^+$ HSPCs were collected and cultured in S50T50 medium under 5% $CO_2$/20% $O_2$ conditions for 2–3 h, and $3 \times 10^5$ HSPCs were transplanted retro-orbitally into lethally-irradiated (8.5Gy using MBR-1520R-3 (Hitachi Power Solutions) with a 125 kV 10 mA, 0.5 mm Al, 0.2 mm Cu filter) recipient Ly5.1 mice noncompetitively.

The sequences of triple-gRNA and the primer set used to confirm KO efficiency were as follows.

sgRNA sequences:

5’-AGACCUGGCUUACCUUUCGU-3’

5’-UGGAGAUGUAAGUCUUACCC-3’

5’-GUUGGUCAGCUUCGGCCCAC-3’

Forward Primer: 5’-CAAAGGAAAAGTCCCATGGAGA-3’

Reverse Primer: 5’-GGGCTTTGGCATGTGGAATG-3’

Sequencing Primer: 5’-CAAAGGAAAAGTCCCATGGAGAATG-3’

For *Pfkfb3* overexpression, HSPCs were cultured in S50T50 medium under 5% $CO_2$/20% $O_2$ conditions for 8–16 h after retroviral transduction, and the equivalent of 3 × $10^5$ HSPCs were noncompetitively transplanted retro-orbitally into lethally-irradiated (8.5Gy using MBR-1520R-3) recipient Ly5.1 mice. After transduction, a group of the cells was cultured in S50T50 medium for 48 h to confirm that transduction (GFP positivity) had been established.

**Cell cycle analysis and apoptosis assay of *Pfkfb3*-KO/overexpressing HSPCs after non-competitive BMT**

BMMNCs were collected from the bilateral femur, tibia, pelvic bone, and sternum of each individual recipient mouse on day 2 after noncompetitive BMT. Recipient BMMNCs were then stained with Lineage-marker-PerCP5.5, Ly5.1-PerCP5.5, and Ly5.2-PE (cell cycle analysis) or Lineage-marker-FITC, Ly5.1-FITC, and Ly5.2-Alexa700 (apoptosis assay). For the analysis, all BMMNCs from each recipient were used in one analysis, and all lineage-marker negative Ly5.2$^+$ cells were analyzed. Cell cycle analysis (Ki67/Hoechst33432 staining) was performed as described in the “**Ki67/Hoechst33432 staining**” section. *In vivo* BrdU labeling assays were performed as reported by Jun et al.[8] using the FITC BrdU Flow Kit (BD Biosciences, Cat# 559619). Apoptosis assays were performed using the PE Annexin V Apoptosis Detection Kit I according to manufacturer instructions.

Cell cycle analysis (Ki67/Hoechst33432 staining) of *Pfkfb3*-overexpressing HSPCs after transplantation was also performed using all BMMNCs from each recipient mouse, and the analysis was performed on all *Pfkfb3*-overexpressing cells (GFP$^+$).

**5-FU administration after BM recovery in *Pfkfb3*-KO HSPCs**

PFKFB3 was gene-edited in HSPCs using triple-gRNA as described above, and the equivalent of 3 × $10^5$ LSK cells were transplanted retro-orbitally into lethally-irradiated (8.5Gy using MBR-1520R-3) recipient Ly5.1 mice noncompetitively. After 2 months, recipient mice were treated with 150 mg/kg of 5-FU intraperitoneally. Peripheral blood was collected on the day of 5-FU administration (day 1), and on days 4, 6, 9, and 16. The dynamics of *Pfkfb3*- or *Rosa*-KO cell abundance (as control group) were analyzed by Sanger sequencing as described above.

**Homing assay of *Pfkfb3*-KO HSPCs**

PFKFB3 was gene-edited in $GFP^+$ HSPCs using triple-gRNA as described above. After editing, $2 \times 10^5$ cells were retro-orbitally transplanted into lethally-irradiated (8.5Gy) C57BL/6 mice. After 16 hours, BMMNCs from recipients were stained for surface antigens and analyzed for the percentage of $GFP^+$ cells within the PI-negative cells.

**Immunocytochemistry**

HSCs from PBS- or 5-FU-treated C57BL/6 mice were subjected to immunocytochemistry using antibodies for PFKFB3 (Abcam, Cat# ab181861), phosphorylated-PFKFB3 (Bioss, Cat# bs-3331R), and methylated-PFKFB3 (developed by Takehiro Yamamoto)[19]. Purified HSCs were resuspended in 50% FCS-PBS and cytospun using the Thermo Scientific Cytospin 4 system (Thermo Fisher Scientific). When using 2-NBDG-positive or -negative HSCs, C57BL/6 mice were given 2-NBDG intravenously (see "***In vivo* 2-NBDG assay**" for details) and subjected to cytospinning. Cytospun cells were fixed using 4% paraformaldehyde in PBS pH 7.4 for 10 min at RT. Fixed cells were washed twice with ice-cold PBS. For permeabilization, cells were incubated for 5 min with PBS containing 0.1% Triton X-100. Permeabilized cells were washed once with ice-cold PBS. After blocking with 3% BSA-PBS for 30 min, cells were incubated in the diluted antibody with 0.3% BSA-PBS in a humidified chamber overnight at 4 °C. A dilution factor of 1:100 was used for all antibodies. The next day, cells were incubated with Goat anti-Mouse IgG2a Secondary Antibody, Alexa Fluor™ 555 (Thermo Fisher Scientific, Cat# A-21137) and DAPI in 0.3% BSA-PBS for 1 h at RT. After two washes with ice-cold PBS, samples were coverslipped with a drop of mounting medium and imaged with a Zeiss LSM 880 microscope (ZEISS, Jena, Germany). Images were acquired at room temperature under darkened conditions using a 100x oil immersion lens. The obtained image data was analyzed using Imaris software (Bitplane) to calculate the MFI of the target for each cell.

**RNA sequencing**

Library preparation for RNA-seq was performed on 3000–3500 HSCs derived from mice after 5-FU or PBS administration. Total RNA was prepared using Rneasy Micro kit (QIAGEN, Hilden, Germany). cDNA was synthesized and amplified using SMART-Seq v4 Ultra Low Input RNA Kit for Sequencing (Takara Bio, Inc., Shiga, Japan). RNA-seq libraries were prepared using the Nextera XT Kit (Illumina,

San Diego, CA, USA). Single-end 75 bp sequencing was performed on a NextSeq 500 platform (Illumina). RNA-seq data were obtained from three independent experiments (biological duplicates) for each cell type. TopHat (version 2.0.13; with default parameters) was used for mapping to the reference genome (UCSC/mm10) with annotation data from iGenomes (Illumina). Then, gene expression levels were quantified using Cuffdiff (Cufflinks version 2.2.1; with default parameters).

**MACSQuant analysis of cell number**

After single GO-ATeam2 knock-in HSC culture, most of the medium (150–170 μL) in wells of a 96-well plate was aspirated and samples were stained with 10 μL antibody cocktail for 30 min at 4 °C. Antibodies used were anti-lineage markers (CD4, CD8a, Gr-1, Mac-1, B220, Ter-119)-PerCP-Cy5.5, anti-c-Kit-APC-Cy7, anti-Sca-1-PE-Cy7, anti-CD150-BV421, and anti-CD48-APC for LSK-SLAM analysis. Antibody cocktail was prepared by mixing 0.1 μL of each antibody. After incubation, 100 μL PBS + 2% FCS was added to wells, and the plates were centrifuged for 5 min at 4 °C and 400 × *g* with low acceleration and medium deceleration. Then, 100 μL supernatant was aspirated and cell pellets were resuspended in 200 μL PBS + 2% FCS + 0.1% PI + 0.25% Flow-Check Fluorospheres (Beckman Coulter, Brea, CA, USA, Cat# A69183). Samples were acquired in fast mode in the MACSquant analysis settings, and volumes of 100 μL (large colonies) or 150–170 μL (small colonies) were analyzed. Data were exported as FCS files and analyzed using FlowJo software. Cell number was corrected by bead count of Flow-Check (~1000 cells/μL). HSCs were counted using $CD150^{+}CD48^{-}$ LSK cell counts. Megakaryocytes were identified as cells with high forward scatter and side scatter, as well as high CD150 and CD41 expression.

**cDNA synthesis and quantitative RT-PCR**

cDNA synthesis and RT-PCR using PFKFB3CA overexpressing cells were performed as previously reported. The primers used were as follows:

MA069663-F: 5′-GGGCATGGCGAGAATGAGTACAA-3′

MA069663-R: 5′-TTCAGCTGGGCTGGTCCACAC-3′

**Statistical analysis**

Data are presented as means ± SD unless otherwise stated. For multiple comparisons, statistical significance was determined by Tukey's multiple comparison test using the Tukey HSD function in the R ×64 4.0.3 software (R Core Team, Vienna, Austria). A paired or unpaired two-tailed Student’s *t*-test and two-way ANOVA with Sidak’s test were used for experiments with two groups. A p-value < 0.05

was considered statistically significant.

| Custom RPMI medium for ATP analysis | | | |
|---|---|---|---|
| **Amino Acids** | **Concentration (mg/L)** | **Vitamins** | **Concentration (mg/L)** |
| L-Arginine HCl | - | D-Biotin | 0.2 |
| L-Asparagine H2O | - | D-Calcium pantothenate | 0.25 |
| L-Aspartic acid | - | Choline chloride | 3 |
| L-Cystine 2HCl | - | Folic acid | 1 |
| L-Glutamic acid | - | i-Inositol | 35 |
| L-Glutamine | - | Niacinamide | 1 |
| Glycine | - | p-Aminobenzonic acid (PABA) | 1 |
| L-Histidine HCl H2O | - | Pyridoxine HCl | 1 |
| Hydroxy-L-proline | - | Riboflavin | 0.2 |
| L-Isoleucine | - | Thiamine HCl | 1 |
| L-Leucine | - | Vitamin B12 | 0.005 |
| L-Lysine HCl | - | **Inorganic salts** | **Concentration (mg/L)** |
| L-Methionine | - | Calcium nitrate (Ca(NO3)2 4H2O) | 100 |
| L-Phenylalanine | - | Potassium chloride (KCl) | 400 |
| L-Proline | - | Magnesium sulfate (MgSO4) | 48.84 |
| L-Serine | - | Sodium chloride (NaCl) | 6000 |
| L-Threonine | - | Sodium bicarbonate (NaHCO3) | 2000 |
| L-Tryptophan | - | Sodium phosphate (Na2HPO4) | 800 |
| L-Tyrosine 2Na 2H2O | - | **Other componemts** | **Concentration (mg/L)** |
| L-Valine | - | Glutathione reduced | 1 |
| **Supplements** | **Concentration (mg/L)** | HEPES | 5960 |
| Sodium lactate | - | Thymidine | 0.3633 |
| BSA | - | Phenol red | - |
| Cholesterol | - | Sodium pyruvate | - |
| Oleic acid | - | **Sugars** | **Concentration (mg/L)** |
| Palmitic acid | - | D-Glucose | - |

**Table S1**. Custom RPMI medium for culture and ATP analysis

Composition of custom RPMI medium for culture (upper) and ATP analysis (lower). "-" means 0 mg/L.

**Table S2**. *In vitro* tracer analysis for 5-FU-treated HSCs (uploaded separately as an Excel file)

Results of tracer analysis using U-$^{13}C_6$-glucose with HSCs from mice treated with PBS or 5-FU. Each section contains raw data from the glycolytic system, TCA cycle, and PPP~NAS from top to bottom. Data from three individual experiments are described for each. All values represent average metabolite levels in single HSCs obtained by dividing the metabolite levels detected in HSCs (compared to internal standards) by the number of HSCs used in the analysis.

**Table S3**. *In vitro* tracer analysis for oligomycin-treated HSCs (uploaded separately as an Excel file)

Results of tracer analysis using U-$^{13}C_6$-glucose with HSCs treated with DMSO (Oligomycin-) or oligomycin (Oligomycin+). Each section contains raw data from the glycolytic system, TCA cycle, and PPP~NAS from top to bottom. Data from four individual experiments are described for each. All values represent average metabolite levels in single HSCs, obtained by dividing the metabolite levels detected in HSCs (compared to internal standards) by the number of HSCs used in the analysis.

**Table S4.** $^{13}C$ quantitative metabolic flux analysis (uploaded separately as an Excel file)
Metabolic flux values of each enzyme obtained from 100 trials of $^{13}C$ quantitative metabolic flux analysis for PBS-treated (left), 5-FU-treated (middle), and OXPHOS-inhibited HSCs (right).

**Table S5.** *In vivo* tracer analysis for 5-FU treated mice (uploaded separately as an Excel file)
Results of tracer analysis during continuous *in vivo* administration of U-$^{13}C_6$-glucose to mice treated with 5-FU or PBS. A sheet is prepared for each metabolite and each contains two tables. The A.U. table (left) shows the metabolite levels detected in the four biological replicates in the 5-FU and PBS groups, obtained by dividing the metabolite levels detected in HSCs (compared to internal standards) by the number of HSCs used in the analysis. The ratio table (right) shows the calculated percentage of labeled metabolites among detected metabolites, where 12C indicates unlabeled metabolites and 13Cn indicates n-carbon labeled metabolites by U-$^{13}C_6$-glucose.

| **GO-ATeam2 genotyping** | |
|---|---|
| Primer name | Sequence |
| 55_CAGGS-5F | 5’-AGAGCCTCTGCTAACCATGTTCATGCCTTC-3’ |
| 570_KusabiraOrange-141R | 5’-GTGACACTAAGTCAAACGCGAAA-3’ |

**Table S6**. Primer list for genotyping PCR

### Figure S1

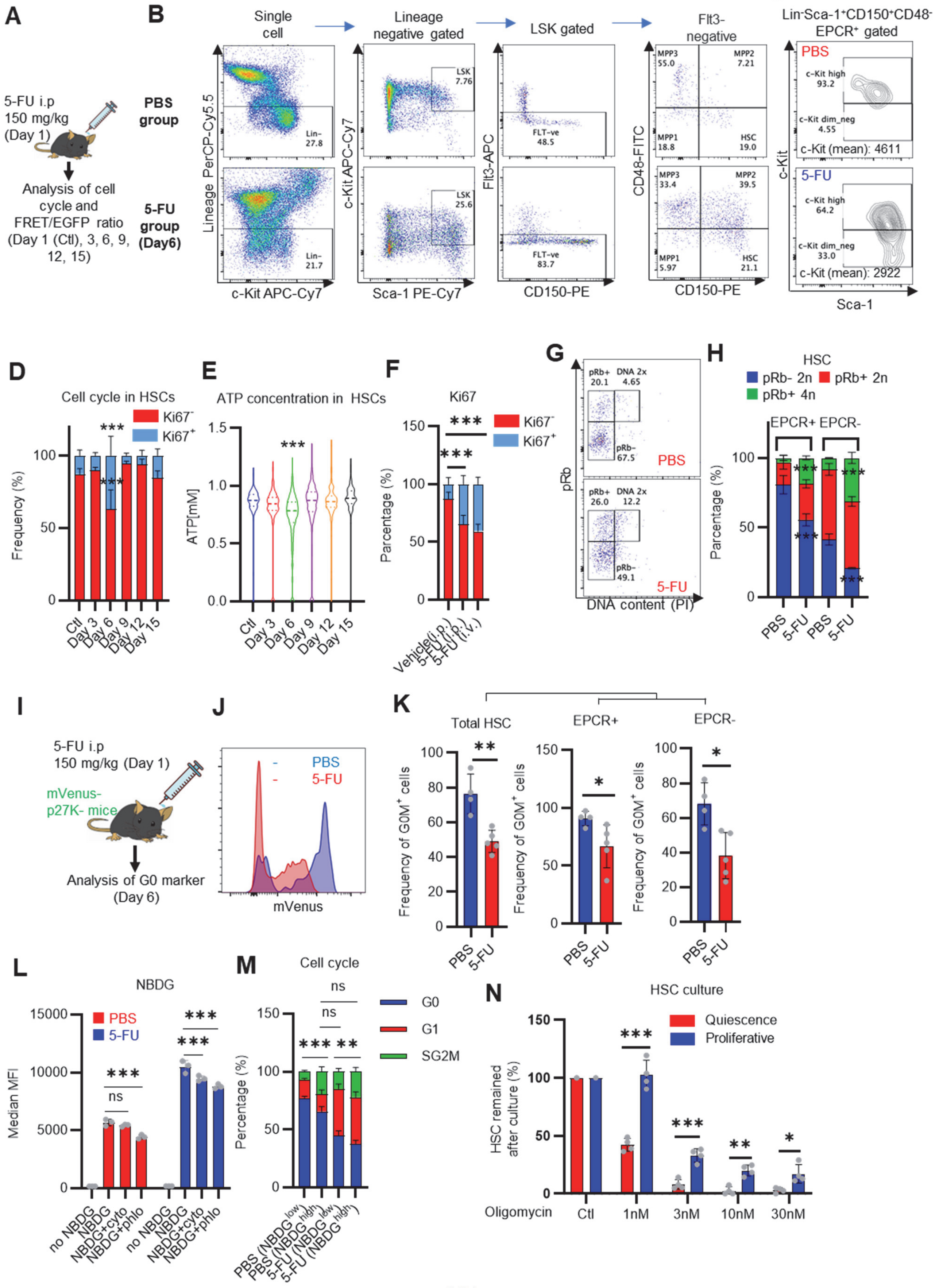
A
5-FU i.p
150 mg/kg
(Day 1)
Analysis of cell cycle and FRET/EGFP ratio (Day 1 (Ctl), 3, 6, 9, 12, 15)
B
PBS group
5-FU group (Day6)
Single cell
Lineage negative gated
LSK gated
Flt3- negative
Lineage PerCP-Cy5.5
c-Kit APC-Cy7
Sca-1 PE-Cy7
Flt3-APC
CD150-PE
CD48-FITC
C
Lin-Sca-1+CD150+CD48- EPCR+ gated
PBS
5-FU
c-Kit
Sca-1
c-Kit (mean): 4611
c-Kit (mean): 2922
D
Cell cycle in HSCs
Frequency (%)
Ki67-
Ki67+
Ctl
Day 3
Day 6
Day 9
Day 12
Day 15
E
ATP concentration in HSCs
ATP[mM]
F
Ki67
Percentage (%)
Vehicle(i.p.)
5-FU (i.p.)
5-FU (i.v.)
G
pRb
DNA content (PI)
H
HSC
pRb- 2n
pRb+ 2n
pRb+ 4n
EPCR+
EPCR-
PBS
5-FU
I
5-FU i.p
150 mg/kg (Day 1)
mVenus-p27K- mice
Analysis of G0 marker (Day 6)
J
mVenus
K
Total HSC
EPCR+
EPCR-
Frequency of G0M+ cells
L
NBDG
Median MFI
no NBDG
NBDG
NBDG+cyto
NBDG+phlo
M
Cell cycle
G0
G1
SG2M
PBS (NBDG low)
PBS (NBDG high)
5-FU (NBDG low)
5-FU (NBDG high)
N
HSC culture
Quiescence
Proliferative
HSC remained after culture (%)
Oligomycin
Ctl
1nM
3nM
10nM
30nM

**Figure S1. Dependence on glycolysis increases with cell cycle progression of HSCs; related to Figure 1**

**(A)** Schematic illustration of 5-FU administration and analysis. **(B)** Representative staining plot of BM cells derived from mice 5 d (day 6) after treatment with PBS or 5-FU (day 1); note the gating of c-Kit high-dim cells in the LSK staining. Because the same BMMNC counts were obtained for each group, the Flt3-negative gated dot plot (bottom-right) for the 5-FU group appears to show an increase in the HSC fraction. (**C**) Contour plot of c-Kit expression in Lin$^-$Sca-1$^+$EPCR$^+$CD150$^+$CD48$^-$ cells from mice five days after PBS or 5-FU administration. **(D)** Frequency of Ki67-positive and -negative HSCs after 5-FU administration. n = 5 biological duplicates for each group. **(E)** Changes in ATP concentration in HSCs after 5-FU administration (n >70 single HSCs for each group). Data are representative results of pooled samples of two biological replicates. **(F)** Ki-67 positivity in HSCs by route of 5-FU administration (i.p. or i.v.). n = 4 biological duplicates for each group. **(G-H)** Intracellular staining of pRb in EPCR$^+$ or EPCR$^-$ HSCs derived from PBS- or 5-FU-treated mice. Representative plot of pRb and DNA content in EPCR$^+$ HSCs from both groups (G). Summary of results (H). n = 3 biological duplicates for each group. **(I-K)** Analysis of mVenus-p27K- mice treated with PBS or 5-FU. Experimental schema (I). Representative G0 marker distribution in HSC in PBS (blue) or 5-FU (red) groups (J). Percentage of G0 marker-positive cells in total HSCs and EPCR$^+$ or EPCR$^-$ HSCs in PBS (blue bars) or 5-FU group (red bars) (K). n = 4–5 biological replicates for each group. The data for each panel is extracted from the same individual. (**L**) *In vitro* 2-NBDG assay. cyto: cytochalasin, phlo: phloretin. n = three biological replicates for each group. (M) *In vivo* administration of 2-NBDG followed by the Ki67/Hoechst 33432 staining of HSCs. n = six biological replicates for each group. **(N)** Relative percentage of HSCs remaining after culture under quiescence-maintaining or proliferative conditions in the presence of oligomycin. HSC number for the control (Ctl) vehicle (DMSO)-treated group was set to 100%; n = 4 technical replicates for each group. The data are representative results from three independent experiments. (See "**MACSQuant analysis of cell number**" under "**Supplementary Methods**" for more information.)

Data are presented as mean ± SD. * $p \leq 0.05$, ** $p \leq 0.01$, *** $p \leq 0.001$ as determined using Student's *t*-test (H, K, N) or one-way ANOVA followed by Tukey's test (D–F, and L–M).

#### Figure S2

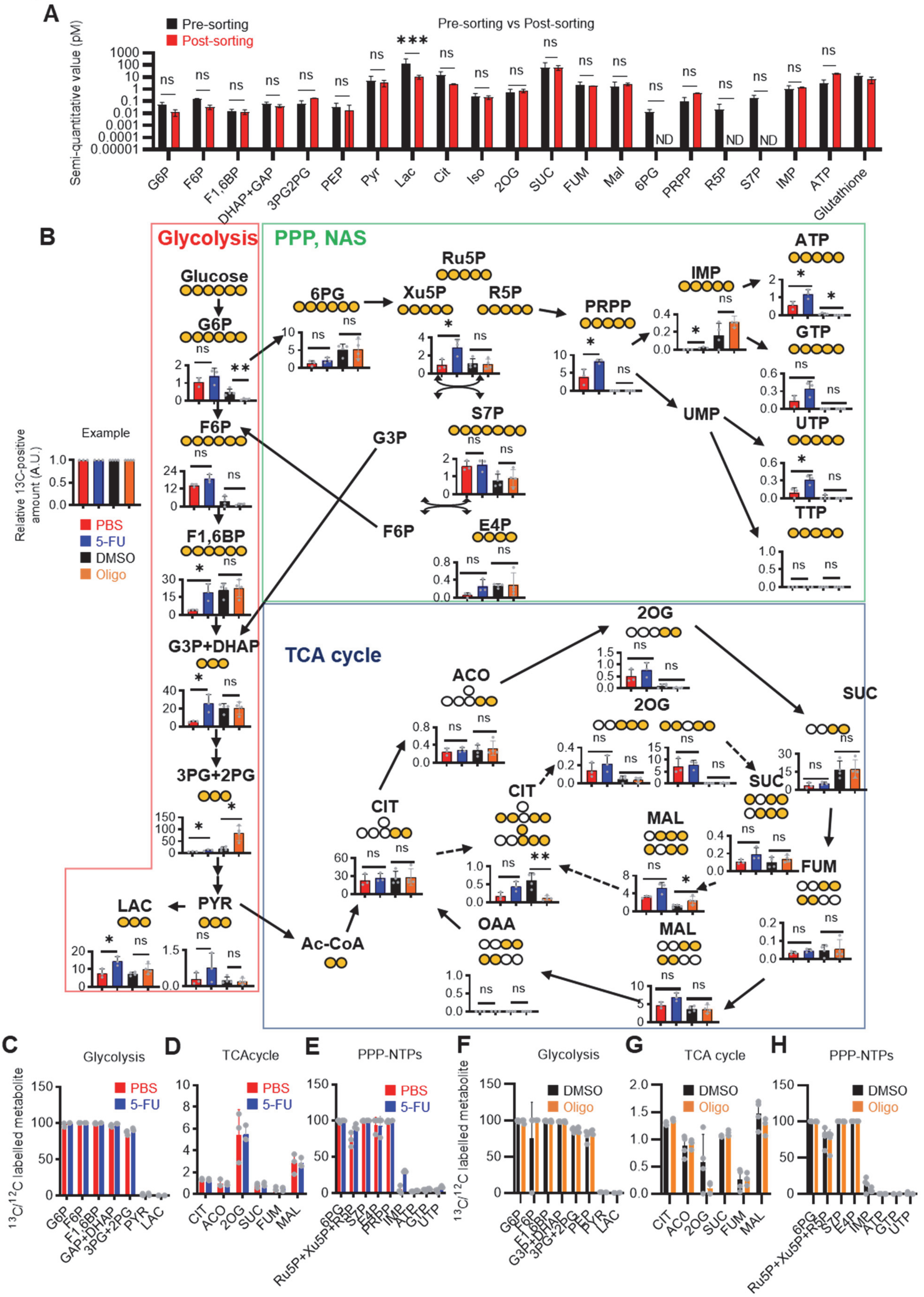

A
Pre-sorting
Post-sorting
Pre-sorting vs Post-sorting
Semi-quantitative value (pM)
G6P F6P F1,6BP DHAP+GAP 3PG2PG PEP Pyr Lac Cit Iso 2OG SUC FUM Mal 6PG PRPP R5P S7P IMP ATP Glutathione
ND
B
Glycolysis
PPP, NAS
TCA cycle
Example
Relative 13C-positive amount (A.U.)
PBS
5-FU
DMSO
Oligo
Glucose G6P F6P F1,6BP G3P+DHAP 3PG+2PG PYR LAC
6PG Ru5P Xu5P R5P PRPP IMP ATP GTP UMP UTP TTP S7P E4P G3P F6P
ACO 2OG SUC CIT FUM MAL OAA Ac-CoA
C Glycolysis
D TCAcycle
E PPP-NTPs
F Glycolysis
G TCA cycle
H PPP-NTPs
$^{13}C/^{12}C$ labelled metabolite

**Figure S2. Quantified metabolite pool in HSCs under quiescence, proliferation, or OXPHOS-inhibition; related to Figure 1, 2**

**(A)** Comparison of metabolite levels in c-Kit enriched cells pre- and post-sorting. n = 3 biological replicates for each group. ND: not detected. **(B)** Metabolic overview of U-13C6-glucose tracing among pathways related to glycolysis, PPP, NAS, and the TCA cycle in HSCs from 5-FU- (blue) or PBS-treated (red) mice, and DMSO- (black) or oligomycin (Oligo)-treated (orange) HSCs. Fates of carbons derived from U-$^{13}C_6$-glucose in each metabolite are shown as yellow circles. Each graph indicates relative amounts of U-$^{13}C_6$-glucose-derived metabolites. **(C-E)** Ratio of U-$^{13}C_6$-glucose-labelled to non-labelled metabolites in glycolysis (C), the first round of the TCA cycle (D), and the PPP plus nucleic acid synthesis (E) in PBS- (red bars) and 5-FU-treated (blue bars) HSCs. **(F-H)** Ratio of U-$^{13}C_6$-glucose-labelled to non-labelled metabolites in glycolysis (F), the first round of the TCA cycle (G), and the PPP plus nucleic acid synthesis (H) in HSCs treated with vehicle (black bars) or oligomycin (orange bars).

In (B-H), data are extracted from three biological replicates for HSCs derived from PBS- or 5-FU-treated mice and from four for HSCs after DMSO or oligomycin treatment. Data are presented as mean ± SD. * $p \leq 0.05$, ** $p \leq 0.01$, *** $p \leq 0.001$ as determined using two-way ANOVA with Sidak's test (A) and Student's *t*-test (B) by comparing PBS and 5-FU groups or DMSO and oligomycin groups.

Abbreviations: G6P, glucose-6-phosphate; F6P, fructose-6-phosphate; F1,6BP, fructose-1,6-bisphosphate; G3P, glycerol-3-phosphate; DHAP, dihydroxyacetone phosphate; 3PG, 3-phosphoglycerate; 2PG, 2-phosphoglycerate; PEP, phosphoenolpyruvate; PYR, pyruvate; LAC, lactate; Ac-CoA; acetyl-CoA; CIT, citrate; ACO, cis-aconitic acid, isocitrate; 2OG, 2-oxoglutarate; SUC, succinate; FUM, fumarate; MAL, malate; OAA, oxaloacetate; 6PG, glucose-6-phosphate; Ru5P, ribulose-5-phosphate; Xu5P, xylulose-5-phosphate; R5P, ribose-5-phosphate; S7P, sedoheptulose-7-phosphate; E4P, erythrose-4-phosphate; PRPP, phosphoribosyl pyrophosphate; IMP, inosine monophosphate; ATP, adenosine triphosphate; GTP, guanine triphosphate; UMP, uridine monophosphate; UTP, uridine triphosphate; TTP, thymidine triphosphate.

### Figure S3

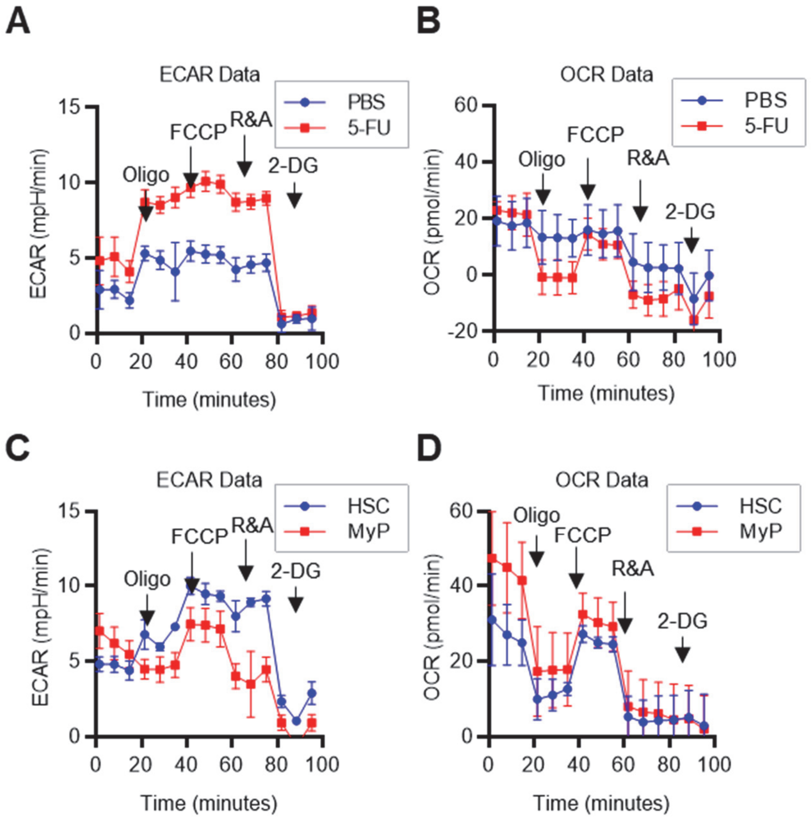


**Figure S3. Mito Stress test results using the Seahorse flux analyzer; related to Figures 1 and 2**
(A–D) Overview of the Mito Stress test on the Seahorse flux analyzer for PBS- or 5-FU-treated HSCs ((A) ECAR, (B) OCR) and HSCs or MyPs ((C) ECAR, (D) OCR).

### Figure S4

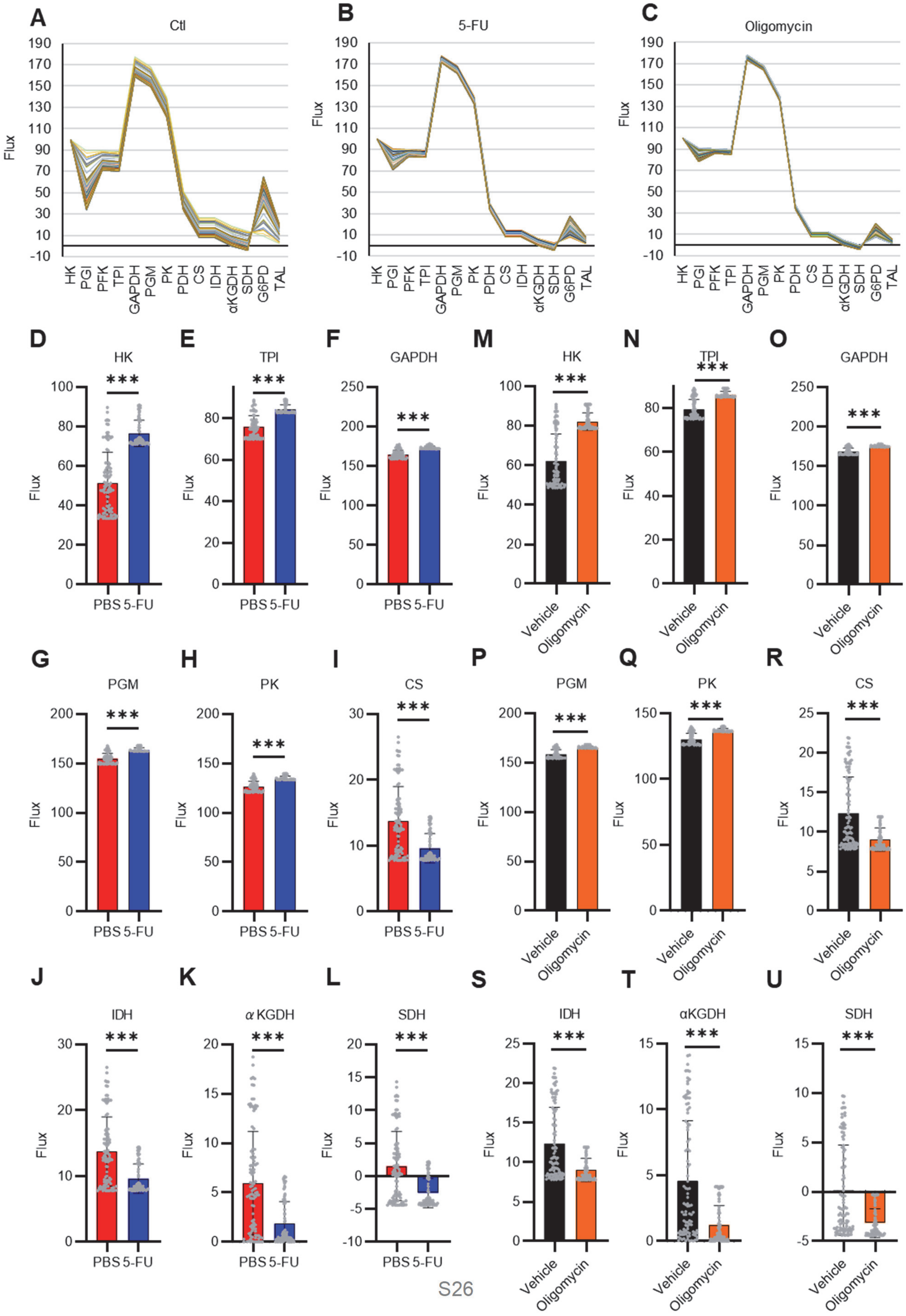
A
Ctl
B
5-FU
C
Oligomycin
Flux
HK
PGI
PFK
TPI
GAPDH
PGM
PK
PDH
CS
IDH
αKGDH
SDH
G6PD
TAL
D
HK
E
TPI
F
GAPDH
M
HK
N
TPI
O
GAPDH
G
PGM
H
PK
I
CS
P
PGM
Q
PK
R
CS
J
IDH
K
α KGDH
L
SDH
S
IDH
T
αKGDH
U
SDH
***
PBS 5-FU
Vehicle
Oligomycin

**Figure S4. Quantitative $^{13}$C-MFA of HSCs under quiescence, proliferation, and OXPHOS inhibition; related to Figure 3**

**(A-C)** Enzyme reaction flux values for each simulation (100 times in total) in PBS-treated (A), 5-FU-treated (B), and OXPHOS-inhibited HSCs (C). Flux values calculated in the same simulation are connected by lines; note the small variation in flux values calculated in different simulations in 5-FU-treated (B) or OXPHOS-inhibited HSCs (C) compared to that in PBS-treated HSCs (A). **(D-U)** Fluxes of each reaction determined using quantitative $^{13}$C-MFA in HSCs from mice treated with 5-FU (blue bars) or PBS (red bars) (D-L), or in HSCs after treatment with vehicle (black bars) or oligomycin (orange bars) (M-U). The net flux was calculated when the glucose uptake was set at 100. The name of the enzyme catalyzing each reaction is listed above the graph. Each gray dot represents the estimated flux obtained from 100 mathematical simulations. (See "**Quantitative $^{13}$C-MFA with OpenMebius**" under "**Supplemental Methods**" for more information.)

Data are presented as mean ± SD. * $p \leq 0.05$, ** $p \leq 0.01$, *** $p \leq 0.001$ as determined using Student's *t*-test (D-U). Abbreviations: HK, hexokinase; PGI, glucose-6-phosphate isomerase; PFK, phosphofructokinase; TPI, triose phosphate isomerase; GAPDH, glyceraldehyde-3-phosphate dehydrogenase; PGM, phosphoglycerate mutase; PK, pyruvate kinase; LDH, lactate dehydrogenase; PC, pyruvate carboxylase; PDH, pyruvate dehydrogenase; CS; citrate synthase; IDH, isocitrate dehydrogenase; αKGDH, α-ketoglutaric acid dehydrogenase; SDH, succinate dehydrogenase; G6PD, glucose-6-phosphate dehydrogenase; TAL, transaldolase.

#### Figure S5

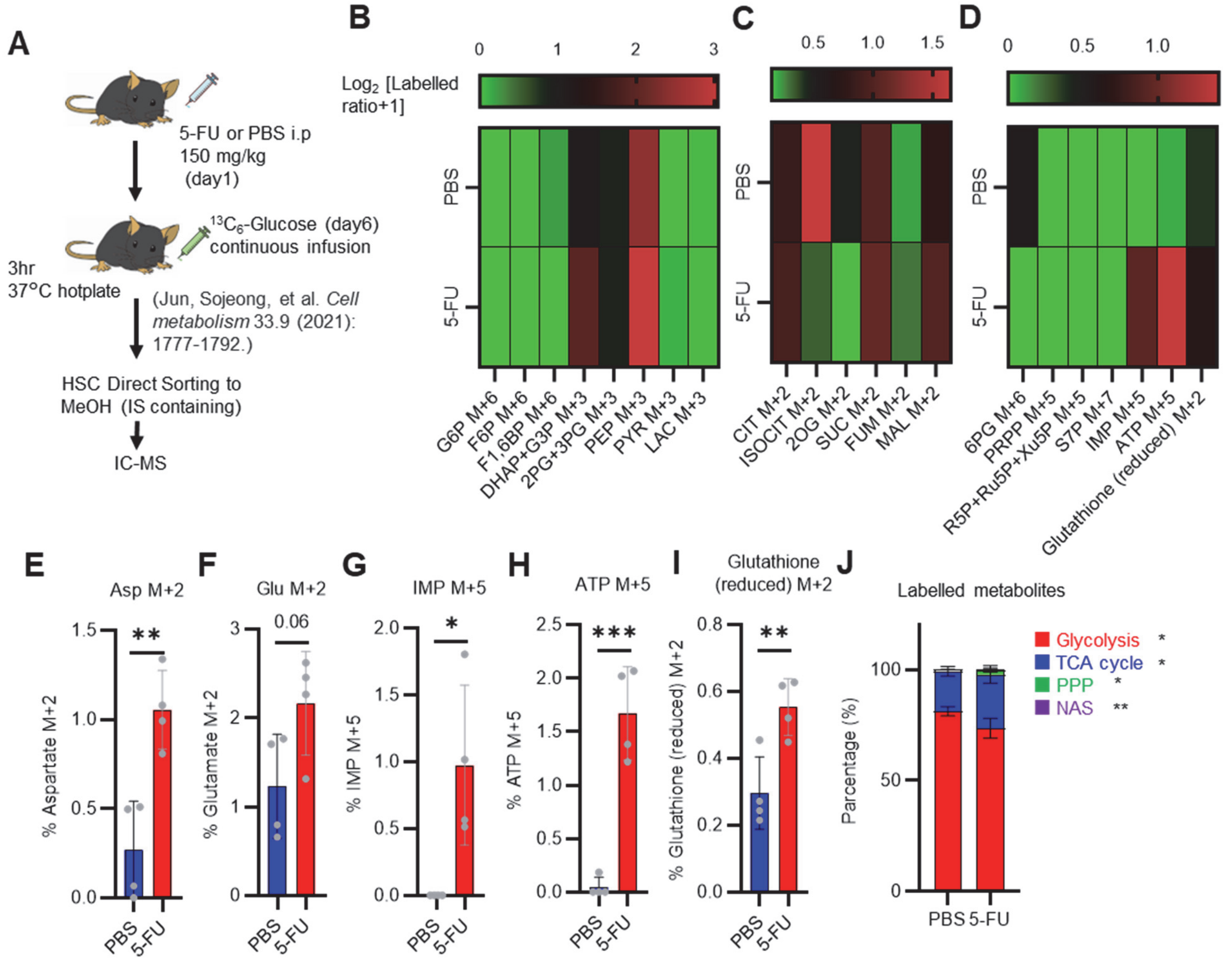


**Figure S5. Quantified metabolite pool in HSCs from PBS- or 5-FU-treated mice**
**(A)** Experimental schema. **(B-D)** Heat maps of the glycolytic system (B), TCA cycle (C), PPP and NAS and glutathione labeling rates (D). **(E-I)** Labeling rates of Asp M+2 (E), Glu M+2 (F), IMP M+5 (G), ATP M+5 (H), and reduced glutathione M+2 (I) in PBS- (blue bars) or 5-FU-treated HSCs (red bars). **(J)** Percentage of total $^{13}$C labeled body mass of glycolysis, TCA cycle, PPP, and NAS detected in PBS-treated or 5-FU-treated HSCs. HSCs derived from one or two mice in the PBS group and two or three mice in the 5-FU group were pooled. n = 4 biological replicates for each group. (See "**Preparation and storage of *in vivo* U-$^{13}C_6$-glucose tracer samples**" in "**Supplementary Methods**" for details.) Data are presented as mean ± SD. * $p \leq 0.05$, ** $p \leq 0.01$, *** $p \leq 0.001$ as determined using Student's *t*-test (E-J).
Abbreviations: G6P, glucose-6-phosphate; F6P, fructose-6-phosphate; F1,6BP, fructose-1,6-bisphosphate; G3P, glycerol-3-phosphate; DHAP, dihydroxyacetone phosphate; 3PG, 3-phosphoglycerate; 2PG, 2-phosphoglycerate; PEP, phosphoenolpyruvate; PYR, pyruvate; LAC, lactate; CIT, citrate; ISOCIT; isocitrate; 2OG, 2-oxoglutarate; SUC, succinate; FUM, fumarate; MAL, malate; 6PG, glucose-6-phosphate; Ru5P, ribulose-5-phosphate; Xu5P, xylulose-5-phosphate; R5P,

ribose-5-phosphate; S7P, sedoheptulose-7-phosphate; PRPP, phosphoribosyl pyrophosphate; IMP, inosine monophosphate; ATP, adenosine triphosphate; Asp, Aspartic acid; Glu, Glutamate.

#### Figure S6

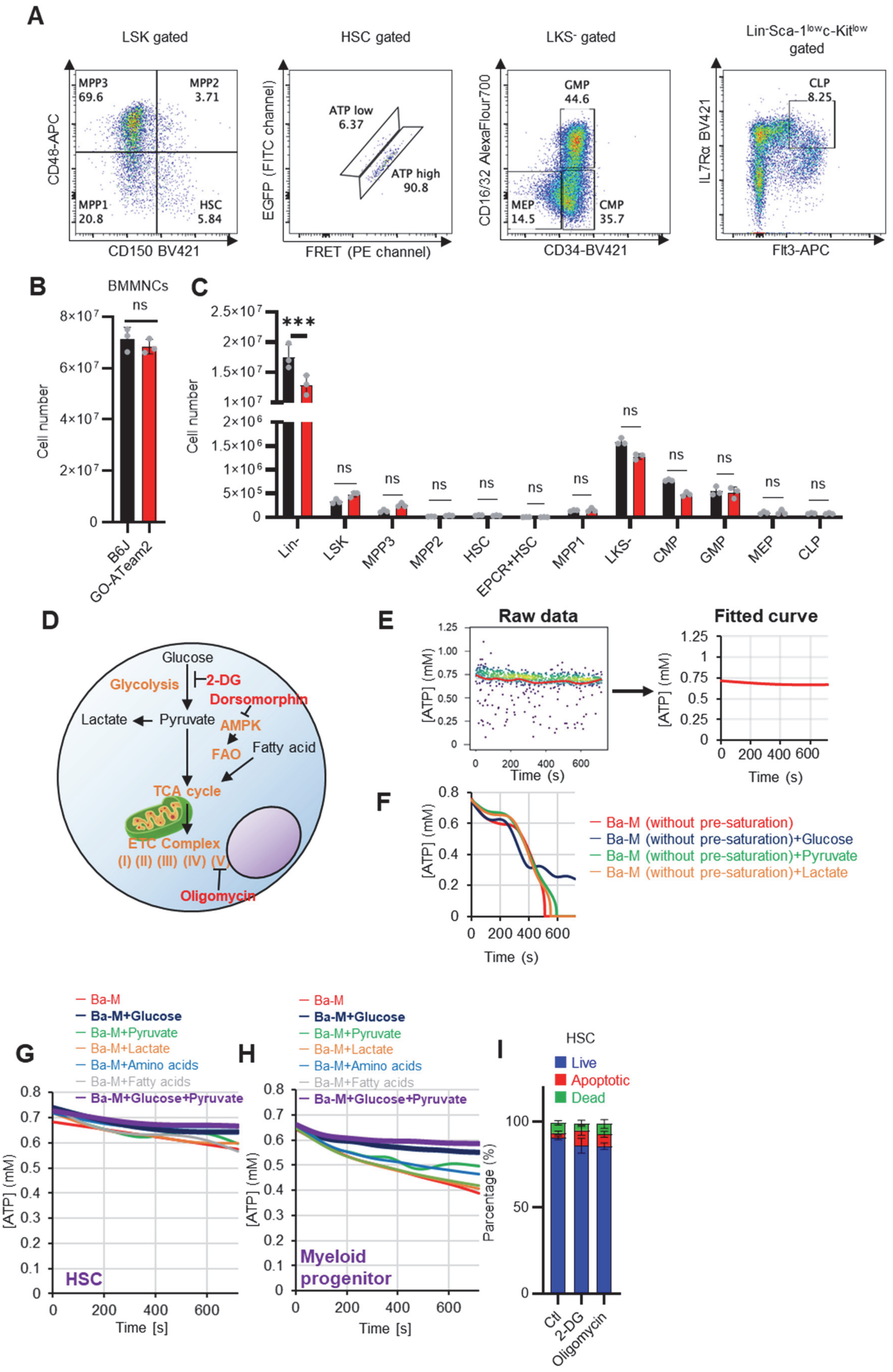
A
LSK gated
HSC gated
LKS- gated
Lin-Sca-1lowc-Kitlow gated
MPP3 69.6
MPP2 3.71
MPP1 20.8
HSC 5.84
CD48-APC
CD150 BV421
ATP low 6.37
ATP high 90.8
EGFP (FITC channel)
FRET (PE channel)
GMP 44.6
MEP 14.5
CMP 35.7
CD16/32 AlexaFlour700
CD34-BV421
CLP 8.25
IL7Rα BV421
Flt3-APC
B
BMMNCs
ns
Cell number
B6J
GO-ATeam2
C
***
ns
Lin-
LSK
MPP3
MPP2
HSC
EPCR+HSC
MPP1
LKS-
CMP
GMP
MEP
CLP
D
Glucose
Glycolysis
2-DG
Dorsomorphin
Lactate
Pyruvate
AMPK
FAO
Fatty acid
TCA cycle
ETC Complex
(I) (II) (III) (IV) (V)
Oligomycin
E
Raw data
Fitted curve
[ATP] (mM)
Time (s)
F
Ba-M (without pre-saturation)
Ba-M (without pre-saturation)+Glucose
Ba-M (without pre-saturation)+Pyruvate
Ba-M (without pre-saturation)+Lactate
G
H
Ba-M
Ba-M+Glucose
Ba-M+Pyruvate
Ba-M+Lactate
Ba-M+Amino acids
Ba-M+Fatty acids
Ba-M+Glucose+Pyruvate
HSC
Myeloid progenitor
Time [s]
[ATP] (mM)
I
HSC
Live
Apoptotic
Dead
Parcentage (%)
Ctl
2-DG
Oligomycin

**Figure S6. Establishment of a real-time ATP concentration analysis system using GO-ATeam2; related to Figure 4**

**(A)** Representative plot of HSPC fractions from GO-ATeam2+ mice. The identified fractions are shown at the top of the graph and the upper gating of that fraction is shown in parentheses. **(B-C)** Number of BMMNCs derived from C57B6/J and GO-ATeam2 mice (B) and percentage of each fraction present (C). **(D)** Schematic diagram showing effects of key metabolic pathways and regulators (orange) and their inhibitors (red). **(E)** Transformation of time-course plot of ATP concentration in individual HSCs from GO-ATeam2 mice (left) to a planar curve after fitting (right). **(F)** Time-course analysis of ATP concentration in HSCs from GO-ATeam2 mice in basal medium (Ba-M) without pre-saturation plus various indicated additives. **(G-H)** Effects of indicated metabolites on ATP concentration of HSCs (G) or MyPs (H) in Ba-M. **(I)** Apoptosis assay results for HSCs exposed to 2-DG (50 mM) or oligomycin (1 μM) for 10 min. n = 3 biological replicates.

Data are presented as mean ± SD. * $p \leq 0.05$, ** $p \leq 0.01$, *** $p \leq 0.001$ as determined using Student's *t*-test (B), two-way ANOVA with Sidak's test (C), or one-way ANOVA followed by Tukey's test (I).

### Figure S7

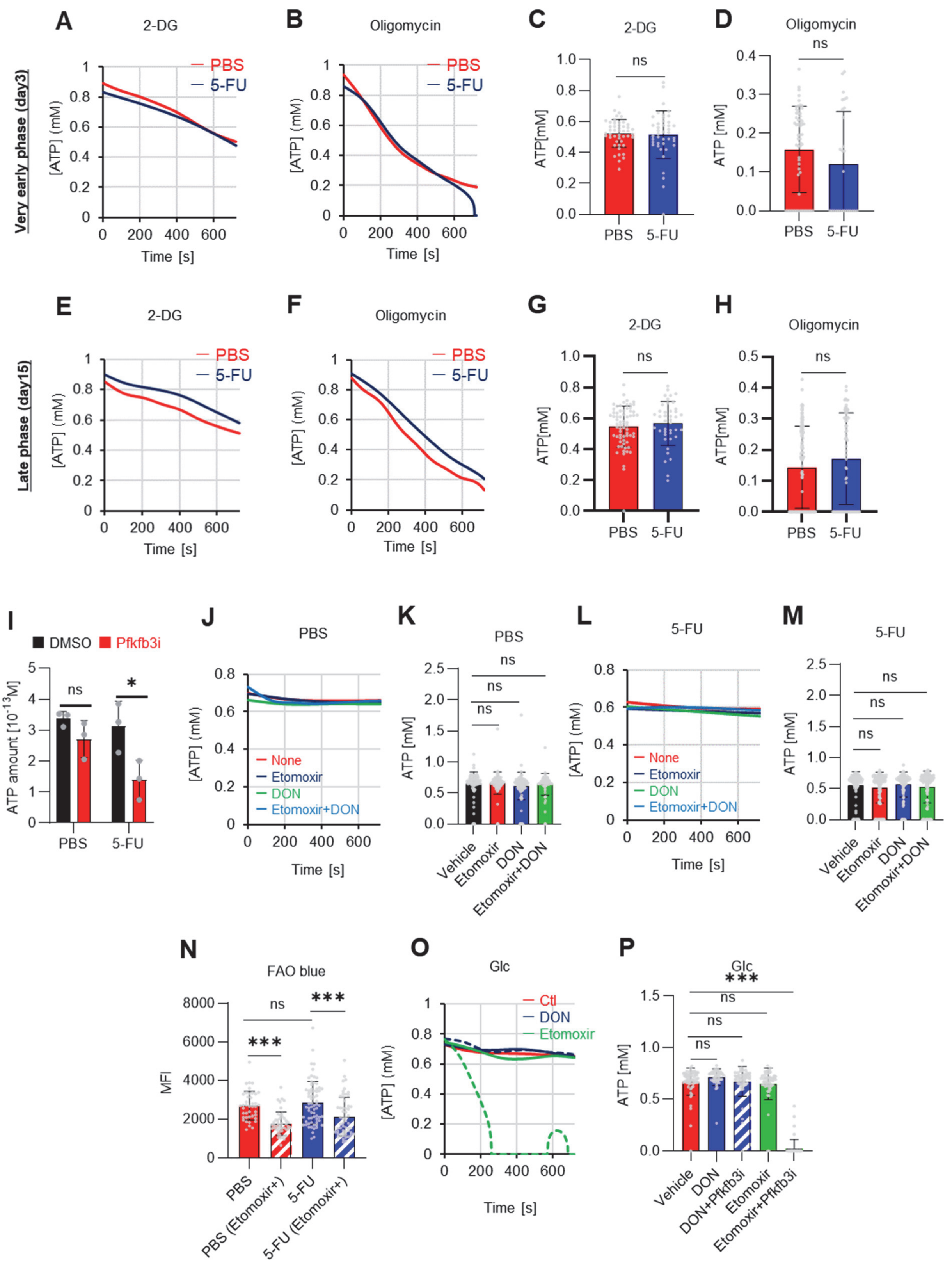


704

705

**Figure S7. FAO is not active in proliferating HSCs; related to Figure 4**

**(A–H)** Results of real-time ATP analysis of PBS- (red) or 5-FU-treated (blue) HSCs (two-days (A–D) or 14 days (E–H) after PBS/5-FU administration) after treatment with 2-DG (A, C, E, G) or oligomycin (B, D, F, H). Bar graphs show corrected ATP concentrations for the last 30 s of the analysis. Each group represents at least 28 cells. Data are representative results of pooled samples from two biological replicates. (see "**Time-course analysis of FRET values**" in "**Supplementary Methods**" for details on the correction method used to calculate the ATP concentration.) (**I**) Average amount of ATP per cell. The amount of ATP detected in the luciferase assay was divided by the number of HSCs used in the analysis. n = three biological replicates for each group. (**J–M**) Results of the real-time ATP analysis of PBS- (J, K) or 5-FU-treated (L, M) HSCs after treatment with etomoxir (100 μM) and/or DON (2 mM). Bar graphs show corrected ATP concentrations for the last 1 min of the analysis. Each group represents at least 60 cells. Data are representative results of pooled samples from two biological replicates. (**N**) MFI of FAOBlue. As a negative control, HSCs were exposed to etomoxir (100 μM). (**O–P**) Effects of DMSO (Ctl, red line), DON (2 mM, blue lines), etomoxir (100 μM, green lines) on ATP in HSCs in the presence of 200 mg/dL glucose. Dashed lines are ATP concentrations with additional AZ PFKFB3 26.

Data are presented as the mean ± SD. * $p \leq 0.05$, ** $p \leq 0.01$, *** $p \leq 0.001$ as determined using a Student's *t*-test (C, D, G, H, and I), or one-way ANOVA followed by Tukey's test (K, M, N, and P).

### Figure S8

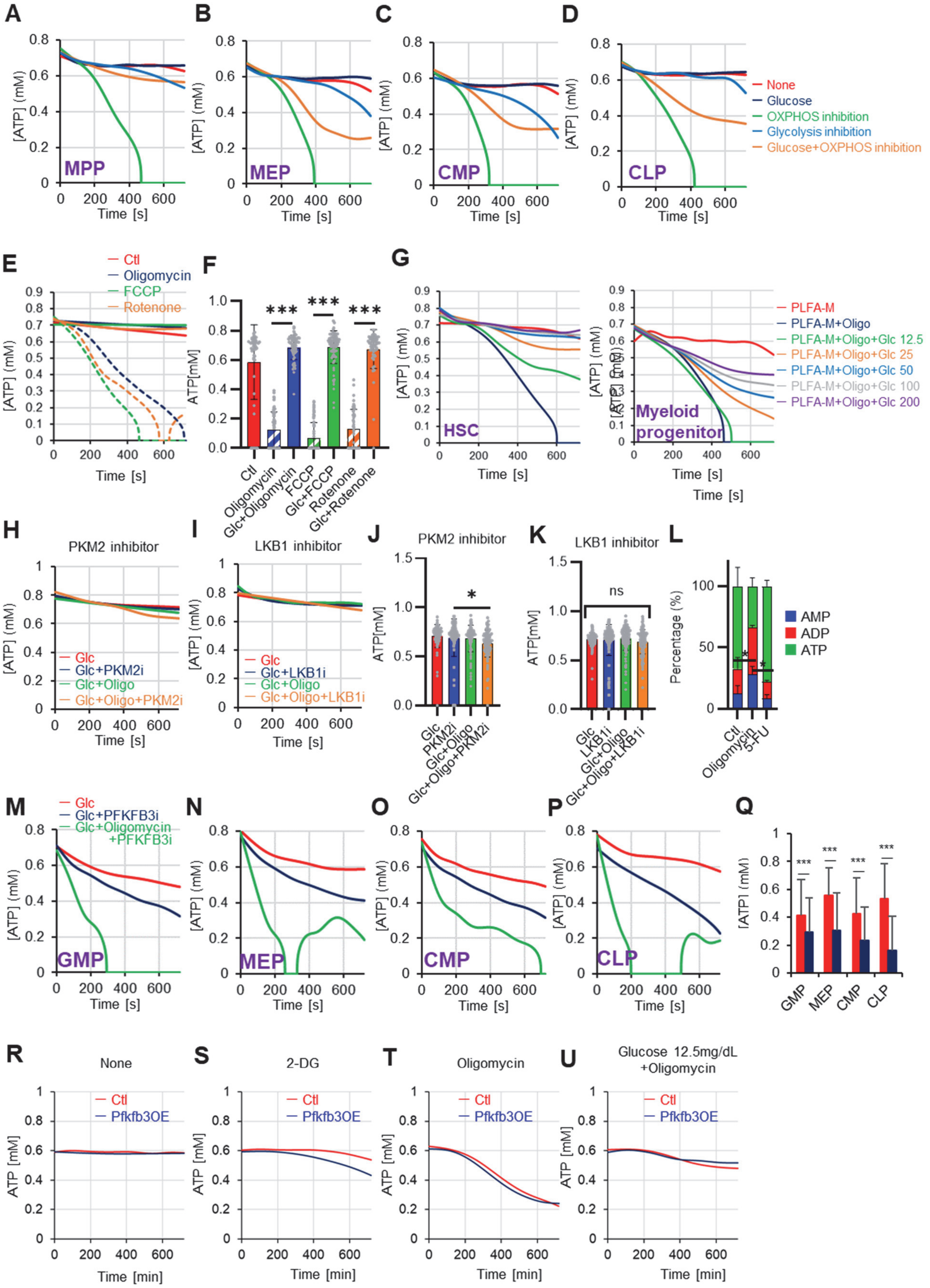


724

**Figure S8. Steady-state Pfkfb3 activity defines HSC and HPC metabolic kinetics and cell cycle; related to Figure 5**

**(A-D)** Evaluation of factors affecting ATP concentration in MPPs (A), MEPs (B), CMPs (C), and CLPs (D) based on the GO-ATeam2 system. GO-ATeam2-knock-in BMMNCs were incubated with glucose, oligomycin, 2-DG, or glucose plus oligomycin, and the FRET/EGFP ratio was calculated. **(E)** Effects of DMSO (Ctl, red line), oligomycin (1 μM, blue lines), FCCP (2 μM, green lines), and rotenone (1 μM, orange lines) on ATP in HSCs. Dashed lines are ATP concentrations with additional dd$H_2O$ and solid lines are ATP concentrations when 200 mg/dL glucose is added. (F) ATP concentration during the last 1 min in (E). Data are representative results for pooled samples of two biological replicates. **(G)** Effects of indicated concentrations of glucose (mg/dL) on ATP concentration in oligomycin-treated or control HSCs (left panel), or in MyPs (right panel) in PLFA medium. **(H-K)** Effects of inhibitors of PKM2 or LKB1 (PKM2i or LKB1i, respectively) on ATP concentration of HSCs from GO-ATeam2 mice in Ba-M with either glucose (Glc) or glucose plus oligomycin. ATP concentrations for the last 2 min of analysis time are summarized in (J) and (K), respectively. Each group represents at least 50 cells. Data are the result of one experiment. **(L)** Composition of adenine phosphates (AMP, ADP, and ATP) in HSCs treated with oligomycin (Oligo), or HSCs from 5-FU-treated mice (5-FU) and control HSCs (Ctl; no treatment or DMSO-treated). Data show results of tracer experiments shown in Figure 1 and Figure 2. **(M-P)** Effects of a PFKFB3 inhibitor (PFKFB3i) on ATP concentration in GMPs (M), MEPs (N), CMPs (O), or CLPs (P) from GO-ATeam2 mice in Ba-M treated with glucose (Glc) plus vehicle (red lines), glucose plus PFKFB3i (blue lines), or glucose plus oligomycin plus PFKFB3i (green lines). **(Q)** ATP concentration in indicated progenitor fractions in Ba-M with vehicle (red bars) or PFKFB3 inhibitor (dark blue bars). ATP concentrations for the last 1 min of the analysis period are shown. Data is summarized from Figure S6 M-P. Each group represents at least 350 cells. Data are representative results of pooled samples from two biological replicates. **(R-U)** Effects of inhibitors on ATP concentration in *Pfkfb3*-overexpressing GO-ATeam2$^+$ HSCs. Cells were exposed to vehicle (Ctl) (R), 2-DG (S), oligomycin (T), or glucose 12.5 mg/dL and oligomycin (U). Data are representative results of pooled samples from two biological replicates.

Data are presented as mean ± SD. * $p \leq 0.05$, ** $p \leq 0.01$, *** $p \leq 0.001$ as determined using Student's *t*-test (F, Q) or one-way ANOVA followed by Tukey's test (J-L).

### Figure S9

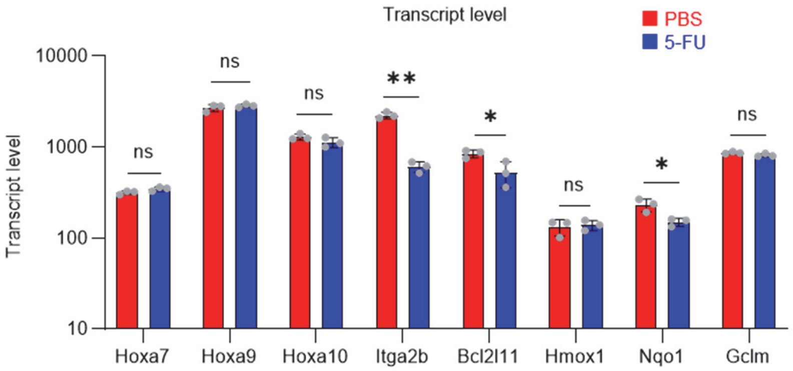


**Figure S9. Gene expression related to Prmt1 in proliferating HSCs; related to Figure 6**
Data are presented as the mean ± SD. * $p \leq 0.05$, ** $p \leq 0.01$, *** $p \leq 0.001$ as determined using a Student's t-test.

#### Figure S10

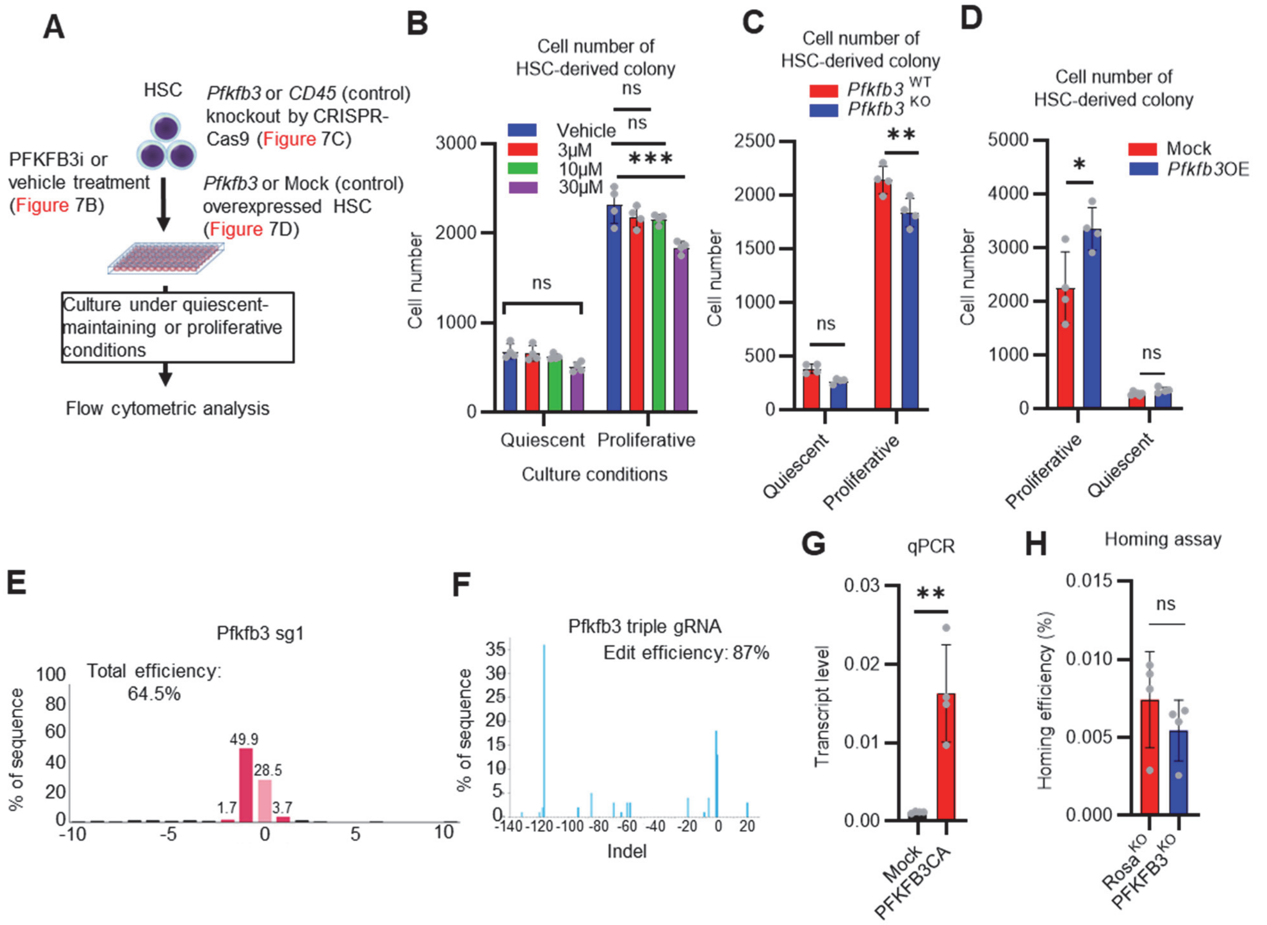


**Figure S10. Pfkfb3 contributes to HSC proliferation and differentiation *in vitro*; related to Figure 7**

(A-D) Effects of *in vitro* PFKFB3 inhibition, KO, or overexpression on HSCs. Experimental design (A). Number of cells in an HSC-derived colony following exposure to a PFKFB3 inhibitor (PFKFB3i) at indicated concentrations (B) or after *Pfkfb3* KO by CRISPR-Cas9 (C); n = 4 technical replicates for each group. Control groups were vehicle (DMSO)-treated (B) or CD45 KO (C). (D) *In vitro* effect of *Pfkfb3*-overexpression on HSCs. Number of cells in *Pfkfb3*-overexpressing or *mock* (pMY-IRES-GFP)-transduced HSC-derived colonies; n = 4 technical replicates for each group. (B-D) are representative results of two or three independent experiments. **(E-F)** KO efficiency evaluated by sgRNA (E) or triple gRNA (F). The indel spectrum (horizontal axis) and the percentage of each indel (vertical axis) are shown. **(G)** qPCR results for *Pfkfb3* expression in validation experiments of *mock*- (black bar) or *PFKFB3CA*- (red bar) overexpression. n = 4 technical replicates for each group. (**H**) Analysis of the homing efficiency and $GFP^+$ blood cells in recipient BM 16 h after the transplantation of Rosa or Pfkfb3 KO $GFP^+$ HSPCs.

Data are presented as mean ± SD. * p ≤ 0.05, ** p ≤ 0.01, *** p ≤ 0.001 as determined using Student's *t*-test (C, D, G, and H) or one-way ANOVA followed by Tukey's test (B).

19. Yamamoto T, Takano N, Ishiwata K, et al. Reduced methylation of PFKFB3 in cancer cells shunts glucose

towards the pentose phosphate pathway. *Nat Commun*. 2014;5:3480.
